# KRAS^G12C^ Inhibition Remodels the Non-Small Cell Lung Cancer Surfaceome to Uncover Targetable Vulnerabilities

**DOI:** 10.64898/2026.10.01.756013

**Authors:** Aparna Padhye, Kalisa Kang, Fan Yi Kong, Nicole Sindoni, Qijia Yu, Baiqi Jiang, Wen-Hsuan Chang, Oluwadara Coker, Eejung Kim, Julien Dilly, Myungsun Shin, Joao A. Paulo, Kathleen Sprouffske, Saskia M. Brachmann, Clint A. Stalnecker, Andrew J. Aguirre, Joseph D. Mancias

## Abstract

Mutant-selective and pan-RAS(ON) multi-selective inhibitors have improved clinical outcomes for patients with RAS-driven cancers; however, adaptive resistance limits the depth and durability of response. Using an integrated genetic, transcriptomic, and mass-spectrometry-based quantitative temporal proteomic platform, we profile the adaptive response to KRAS^G12C^ inhibitors (KRASi) in KRAS^G12C^-mutant lung adenocarcinoma models. Our analyses reveal remodeling of signaling pathways and the cell surface proteome, including cell surface proteins selectively upregulated across cell lines, tumor models, and treatment durations. Folate receptor alpha (FOLR1) emerged as a therapeutically actionable target that is significantly upregulated with KRASi and is part of a broader acute cellular state transition characterized by acquisition of an epithelial identity. Combining KRASi with mirvetuximab soravtansine-gynx, a clinically approved FOLR1-targeting antibody drug conjugate, enhanced tumor cell cytotoxicity and deepened response and tumor control in cellular and in vivo models. Additionally, this work represents a comprehensive multi-omic resource characterizing non-genetic temporal adaptation to KRASi for future mechanistic and therapeutic investigations.

## INTRODUCTION

KRAS^G12C^ mutations account for ∼12% of actionable mutations in non-small cell lung cancer (NSCLC)^1^. The KRAS^G12C^ inhibitors sotorasib and adagrasib have demonstrated meaningful clinical benefit and are now approved by the Food and Drug Administration (FDA) for use in previously treated patients with NSCLC. However, complete and sustained responses are rare, and acquired resistance and clinical progression inevitably occur^2–4^. Resistance is broadly categorized into genetic and non-genetic mechanisms. Genetic resistance includes acquired secondary KRAS mutations that interfere with inhibitor binding within the switch-II pocket^5^ or reactivation of RAS-MAPK signaling via acquired alterations in *NRAS*, *EGFR*, *FGFR2*, and *MYC* which have been identified in a substantial proportion of patients at progression^6,7^. Combinations with targeted therapies have been evaluated preclinically to address these resistance mechanisms but are also expected to eventually succumb to resistance. Non-genetic adaptive mechanisms are increasingly recognized as major drivers of treatment failure. Epithelial-to-mesenchymal transition (EMT) and histological transformation from adenocarcinoma to squamous cell carcinoma (adeno-to-squamous transition, AST) have been observed in patients with acquired resistance to KRASi^8,9^. Additionally, KRASi promotes transition to a quiescent alveolar type 1 (AT1)-like cancer cell state^10^ or a regenerative mucinous transcriptional state that support long-term tumor persistence under KRASi pressure^11^.

Standard-of-care first-line therapy for patients with advanced NSCLC harboring KRAS^G12C^ mutations includes immune checkpoint inhibitors (ICIs) either as monotherapy or in combination with platinum doublet chemotherapy^12,13^. Strategies combining KRAS^G12C^ inhibitors with platinum doublet chemotherapy have been explored to improve upon standard first-line outcomes in lung cancer patients. In the CodeBreaK 101 phase 1b trial, sotorasib plus carboplatin/pemetrexed yielded a first-line objective response rate (ORR) of 65% and median progression free survival (PFS) of 10.8 months. However, grade 3-4 treatment related adverse events (TRAEs) occurred in 52% of patients, with 9% of patients discontinuing sotorasib due to TRAEs^14^. In parallel, early-phase clinical data investigating KRAS^G12C^ inhibition in combination with ICIs have demonstrated preliminary antitumor activity but also highlight important safety considerations, including treatment-related hepatotoxicity and immune-mediated adverse events^15^. Ongoing clinical trials, including KRAScendo-170 (NCT05789082)^16^, are evaluating the efficacy of a KRAS^G12C^ inhibitor with ICI and chemotherapy. The clinical feasibility of such triplet strategies remains uncertain due to potential overlapping toxicities. An alternative strategy for delivering chemotherapy in a targeted manner is using antibody-drug conjugates (ADCs). ADCs deliver cytotoxic payloads directly to tumor cells expressing specific surface antigens while minimizing adverse effects on normal tissues. ADCs have emerged as an important class of anticancer therapies with multiple approvals by the Food and Drug Administration (FDA) to date^17^. Oncogenic KRAS signaling remodels the surfaceome^18^, and recent studies have shown that MAPK pathway inhibition can dynamically alter surface protein expression, potentially increasing the accessibility to antibody-based treatment modalities^19,20^. However, the landscape of the NSCLC surfaceome in the context of KRASi remains incompletely characterized.

Here, we systematically define adaptations to KRASi beyond genomic alterations to understand how tumor cells adapt and survive under KRASi pressure. We perform integrated multi-omic profiling to identify cell surface proteins that are significantly upregulated by KRASi and could represent targets for combination therapy. We identify a global reprogramming of the cell surfaceome upon KRASi across models over different time points. Folate receptor alpha (FOLR1, FRα), a glycosylphosphatidylinositol-anchored receptor involved in folate uptake, emerged as a robustly and reproducibly upregulated cell surface protein upon KRASi^21^.

FOLR1 has limited expression within normal tissues and is overexpressed in malignant tumors of epithelial origin, including ovarian, lung, and breast cancer^22–24^. A FOLR1 targeting ADC, mirvetuximab soravtansine-gynx (Elahere), has been approved by the FDA for use in patients with platinum-resistant epithelial ovarian, fallopian tube, or primary peritoneal cancers^25–28^. Next-generation FOLR1 ADCs are being developed to expand targeting across a broader range of FOLR1 expression levels and to optimize therapeutic index^29^. Several FOLR1-targeting ADCs are also entering clinical evaluation in lung cancer, including luveltamab tazevibulin (REFRαME-L1)^30,31^, MORAb-202^32^ and rinatabart sesutecan (NCT07288177). Collectively, these studies highlight the growing clinical interest in exploiting FOLR1 as a therapeutic target in lung cancer.

Our results show that FOLR1 is an actionable therapeutic target in the context of KRASi and combination of KRASi with FOLR1-ADC enhanced overall efficacy in in vitro and in vivo models and can be utilized for KRAS-mutant FOLR1-positive cancers. Additionally, we provide an unbiased multi-omics dataset to nominate other targets that could be used for combination targeting with KRASi.

## RESULTS

### Multi-omics reveals a temporal adaptive response to KRAS^G12C^ inhibition

KRAS^G12C^ mutant NSCLC tumors have varying responses to KRAS^G12C^ inhibitors (KRASi), reflecting substantial biological heterogeneity. To systematically define both genetic and non-genetic mechanisms of adaptation, we implemented a multi-omics strategy to characterize temporal responses to KRASi across in vitro and in vivo NSCLC models. We profiled the whole exome, transcriptome and proteome of KRAS^G12C^ mutant NSCLC (adenocarcinoma) models treated with an investigational, covalent KRAS^G12C^ inhibitor, JDQ443^33^, at acute and long-term timepoints (Fig. 1A). Acute treatment (2 days) resulted in suppression of pERK which rebounded after 7 days (Supplementary Fig. S1A). To model acquired resistance, we generated isogenic pairs of resistant cell lines cultured in increasing dose of JDQ443 over 3-4 months (Supplementary Fig. S1B-C). Consistent with prior reports^8^, resistant cells underwent epithelial-to-mesenchymal transition (EMT), as determined by increased expression of the mesenchymal transcription factor ZEB1 and concomitant loss of the epithelial adhesion and polarity molecule, E-cadherin (Supplementary Fig. S1D). Additionally, we utilized a cell line derived xenograft model treated with JDQ443, with tumors collected at day 5, 14 and 45 (Fig. 1A, Supplementary Fig. S1E).

**Figure 1.**
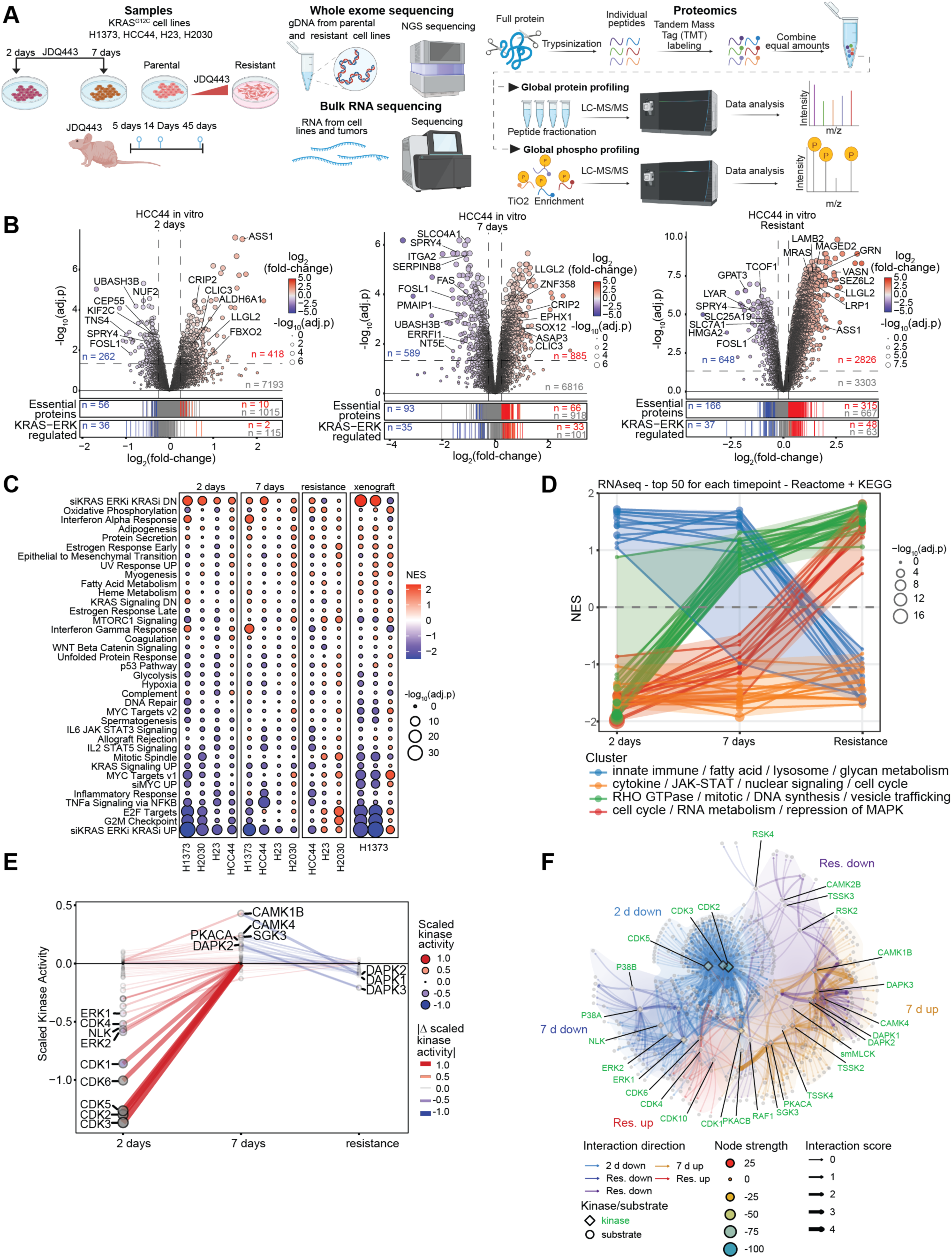
Multi-omic profiling of KRAS^G12C^ lung adenocarcinoma cancer models. **A.** Schematic of data collection for multi-omic profiling for KRAS^G12C^ mutant human lung cancer cell lines and xenograft models treated with JDQ443 at different timepoints. Samples were analyzed by whole exome sequencing, bulk-RNAseq, and multiplexed tandem mass tag (TMT) quantitative global proteomics and phosphoproteomics. **B.** Volcano plots illustrate statistically significant protein abundance differences in HCC44 cells treated with JDQ443 for 2 days, 7 days and to the point of resistance. Volcano plots display the -log_10_ (p value adjusted) versus the median log_2_ fold-change of the relative protein abundance of KRASi-treated to control (DMSO) samples. The number of statistically significant upregulated (red, adjusted p value < 0.05, log_2_(fold-change) > 0.25), downregulated (blue, adjusted p value < 0.05, log_2_(fold-change) < -0.25), or not significant (grey, adjusted p value > 0.05) proteins are shown. Essential proteins (Chronos < -0.75) and proteins within KRAS-ERK regulated gene sets are shown in the bar plot (below). Significantly upregulated (red, adjusted p value < 0.05, log_2_(fold-change) > 0.25), downregulated (blue, adjusted p value < 0.05, log_2_(fold-change) < -0.25), or not significant (grey, adjusted p value > 0.05) are annotated. **C.** GSEA of differential protein expression from all in vitro and in vivo models for Hallmark signatures and KRAS/ERK^37^- or MYC^105^– dependent signatures. **D.** Top 50 differentially enriched or depleted KEGG or Reactome gene sets at each time point from GSEA result of differential transcript expression from all in vitro models combined. Pathways were clustered by temporal pattern of enrichment and general biological pathways determined by Szymkiewicz-Simpson similarity of leading edge genes as shown in Supplemental Fig. S4B. **E.** Relative kinase activities calculated by motif enrichment of phosphoproteomics data and scaled by the product of log_2_(frequency factor) and min-max scaled false discovery rate. Points are filled and sized by the scaled kinase activity and lines are colored and sized by the change between the two connecting timepoints. **G.** Kinase-substrate networks at 2 days, 7 days, or at resistance to KRASi constructed as described previously^40^ using motif preferences and databases available on PhosphoSitePlus and BioGRID. Nodes are sized and colored by their strength in network, and shape represents kinase (diamond) or substrate. Edges are colored by direction of association with up- or down-regulated phosphosites. Edge width represents the magnitude of interaction score. The top quantile of kinases is labeled in each network (green).

We first examined genomic changes associated with resistance by profiling variant allele frequency (VAF) trajectories of selected protein-altering small variants (SNVs and indels) together with genome-wide copy-number alterations across evaluable isogenic cell-line pairs (Supplementary Fig. S1F, G). NCI-H23 showed a small-variant shift, with a slight increase in the VAF of *KRAS p.Q61K* accompanied by a slight decrease in *KRAS p.G12C*, whereas variants in *KEAP1*, *STK11*, and *TP53* remained detectable at broadly similar VAFs (Supplementary Fig. S1F). In contrast, NCI-H2030 showed minimal to no SNV changes among the displayed cancer-relevant variants, with *KRAS p.G12C* and *STK11 p.E317\** remaining stable between vehicle and resistant states. Likewise, HCC44 showed no appreciable VAF change among the displayed variants in *TP53*, *STK11*, *RB1*, *KRAS*, and *KEAP1*. Genome-wide CNVkit profiles showed no recurrent copy-number alteration across models, and most model-specific differences were modest relative to noise, except a possible 17p deletion in *TP53* in NCI-H2030 (Supplementary Fig. S1G). Together, these data do not identify a shared small-variant or copy-number mechanism of acquired KRAS inhibitor resistance in these isogenic cell line models.

While we did not observe second-site mutations in KRAS within our isogenic resistance models, we sought to define the landscape of possible secondary mutations in KRAS^G12C^ that can confer resistance to switch II pocket KRAS^G12C^ inhibitors. To do so, we performed deep mutational scanning^8,34,35^ experiments with two KRAS^G12C^ inhibitors, JDQ443 and sotorasib. Using a lentiviral library that encoded nearly all possible alleles with single amino acid substitution within the *KRAS^G12C^* backbone allele, we performed a positive-selection screen in the Ba/F3 cell line for mutations that cause resistance to sotorasib and JDQ443^8,36^. Drug-resistance mutations were defined as those that enabled cell survival during treatment with a KRAS^G12C^ inhibitor (Supplementary Fig. S2A). Multiple mutations that were identified in the screen at codons 8, 12, 68, 69 and 117 conferred strong resistance to JDQ443 (Supplementary Fig. S2B-C). Structural mapping of residue-level enrichment scores demonstrated that resistance-associated substitutions cluster within and surrounding the Switch II pocket (Supplementary Fig. S2D). In the sotorasib screen, numerous strong resistance mutations were observed at codons 8, 9, 12, 15, 59, 96, and 117, as previously observed^8^ (Supplementary Fig. S2E). A comparison of resistance patterns between JDQ443 and sotorasib showed generally strong concordance (Supplementary Fig. S2F) consistent with the similar mode of KRAS^G12C^ binding and inhibition.

Due to the lack of a convergent genetic mechanism of resistance, we investigated whether adaptive responses instead converge at the proteomic and transcriptomic levels. We found global proteome and transcriptome reprogramming across acute and long-term KRASi. At the proteome level, we observed substantial differential regulation of proteins at 2 days, 7 days, and upon resistance in HCC44 cells, rather than a simple adaptive response that returned to baseline (Fig. 1B, Supplementary Table S1-2). This global proteome reprogramming was also observed in vivo, with substantial dysregulation at 5, 14, and 45 days of treatment (Supplementary Fig. S3A). Correlation analyses showed highest correlations for acute 2-day or 7-day treatments across cell lines that diverged from resistant samples (Supplementary Fig. S3B). The more mesenchymal cell lines HCC44 and H2030 adapted faster to KRASi, where the 7-day treatment clustered with resistant variants. Principal component analysis showed a similar trend, where the more epithelial cell line H23 was separated upon resistance compared to 2-day and 7-day treatment, whereas the mesenchymal cell lines were tightly clustered across time points. At the transcriptome level these trends were similar, except correlations were highest within a given cell line, reflecting the higher coverage of bulk RNAseq and cell line heterogeneity (Supplementary Fig. S3C, Supplementary Table S3). Unlike RNAseq, the coverage of proteins detected by LC-MS proteomics techniques can be highly variable across cell lines and mass-tag sets. We found a high proportion of proteins detected in our datasets were detected across cell lines and time points, with lower proportions only shared with a limited number of cell lines or were cell-line specific (Supplementary Fig. S3E). Additionally, we found high agreement across cell lines for the top up- and down-regulated proteins despite gaps in detection (Supplementary Fig. S3F).

To better understand the adaptation to KRASi, we annotated proteins with high dependence to support proliferation in KRAS-mutant NSCLC cells from DepMap and KRAS-ERK-regulated proteins^37^ (Fig. 1B). We found strong suppression of KRAS-ERK-regulated and essential proteins at short time points both in vitro (2 days) and in vivo (5 days), with few upregulated. This suppression of KRAS-ERK-regulated and essential proteins began to rebound or was maintained at the longer 7-day in vitro and 14-day in vivo treatment, with more essential proteins upregulated compared to acute treatment. However, JDQ443-resistant cells or 45-day in vivo treatment showed a rebound in KRAS-ERK-regulated proteins and a substantial increase of essential proteins in vitro or attenuation of downregulated essential proteins in vivo, consistent with a rebound of activities restoring proliferative capacity of cells and tumors (Supplementary Fig. S3D).

We next performed gene set enrichment analyses (GSEA) on both the proteome and transcriptome across cell lines to identify pathways driving acute versus long-term adaptation (Supplementary Table S4-5). Pathway level changes in the proteome were broadly conserved across conditions despite heterogeneity in individual protein expression (Fig. 1C). We found strong suppression of KRAS-ERK-regulated genes at acute 2-day KRASi, consistent with on-target inhibition. Additionally, we found suppression of the growth regulatory pathways, E2F targets and G2M checkpoint, and enrichment of KRAS-ERK-suppressed proteins (siKRAS ERKi KRASi DN), suggesting strong KRAS inhibition and growth suppression. By 7 days, we observed attenuation of these KRAS-dependent and growth regulatory pathways that approached baseline by the time of resistance. Resistant cells did not show any remarkable patterns of pathway regulation using Hallmark and KRAS-ERK-regulated pathways. This trend of strong KRAS-ERK suppression at shorter time points that rebounded with sustained inhibition was also observed in vivo. We extended our analysis to Reactome and KEGG pathways and observed similar results, where cell cycle-related pathways were suppressed at acute timepoints and rebounded at the point of resistance (Supplementary Fig. S4A). At the transcriptome level, we observed four distinct clusters of pathway regulation across time points (Fig. 1D). As many of these pathways share common genes, we used overlap coefficients determined by the Szymkiewicz-Simpson method using the leading-edge genes to identify redundant pathway terms (Supplementary Fig. S4B). We found gene sets related to cytokine signaling, JAK-STAT, nuclear signaling, and some cell cycle regulatory pathways were suppressed across all timepoints. Two pathway clusters were suppressed that rebounded by 7 days or resistance. Pathways that rebounded by 7 days were related to cytoskeletal signaling, mitosis, DNA synthesis, and vesicle trafficking. Pathways that rebounded by resistance included core cell cycle pathways, RNA metabolism, and suppression of MAPK activity. Together, these trajectories show a step-wise rebound of core cellular machinery required to regain proliferation, including nucleotide biosynthesis and rearrangement of the cellular architecture. Lastly, a set of pathways enriched with acute inhibition were suppressed at the point of resistance. These pathways included innate immune signaling, metabolic pathways such as synthesis of fatty acids, glycans, and lysosomal activities.

To understand the signaling dynamics underlying proteomic adaptation, we next examined the phosphoproteome across treatment timepoints. Like the reprogramming of the proteome and transcriptome, we found global reprogramming of the phosphoproteome (Supplementary Fig. S5A, Supplementary Table S6). Because many phosphosites detected by phosphoproteomics do not have an ascribed function, we annotated phosphosites with known regulatory function^38^ and found broad differential regulation of regulatory sites across time points (Supplementary Fig. S5B). We next used comprehensive phospho-motif profiling^39^ to determine kinase activities associated with the kinome reprogramming. At acute 2-day treatment, we found strong suppression of cyclin-dependent kinases (CDKs) (Fig. 1E, Supplementary Fig. S5C Supplementary Table S7). By 7 days, CDK activity returned to baseline and several upregulated kinases were detected including SGK3, PKN1, CAMK4, PKACA, CAMK1B, and DAPK2. Both downregulated and upregulated kinases were detected in resistant cells, with the top downregulated kinases being DAPK1-3, CAMK, and RSK2, and upregulated kinases CDK4/6 and CDK10. To understand the substrate networks these kinases were regulating, we applied a kinase-substrate network analysis^40^. This analysis revealed a core network of downregulated substrates of ERK and CDKs at 2 days (Fig. 1F, blue edges), followed by a largely separate network of upregulated substrates by 7 days (orange edges). Interestingly, many of the substrates with upregulated phosphosites at 7 days were shared by the downregulated substrates in resistant cells, suggesting these substrates undergo a complex and dynamic regulation by kinome rewiring. Lastly, the network of substrates with upregulated phosphosites in resistant cells (red edges) shared overlap with the 2-day and 7-day downregulated networks, showing the rebound of kinase activities that were initially suppressed by KRASi.

### KRASi induces remodeling of the cellular surfaceome that can be targeted with antibody-drug conjugates

Our global proteomics and transcriptomics showed a systems-wide reprogramming of the cell transcriptome, proteome, and phosphoproteome with acute and sustained KRASi. Consistent with reprogramming of the cell surface, we observed an acute upregulation of canonical epithelial surface markers including CDH1, OCLN, MUC1 alongside suppression of mesenchymal gene sets (AXL, CDH2, VIM, ZEB1) upon KRASi (Fig. 2A). However, the breadth and dynamics of surfaceome remodeling upon KRASi have not been systematically characterized. Given that cell surface proteins can both contribute to adaptive resistance and serve as targets for surface directed therapeutics, including antibody drug conjugates (ADCs), we sought to comprehensively define the surfaceome of KRAS^G12C^ mutant lung cancer cells under KRASi. We utilized a biotinylation based enrichment strategy coupled with mass spectrometry to selectively identify and quantify cell surface accessible proteins (Fig. 2B).

**Figure 2.**
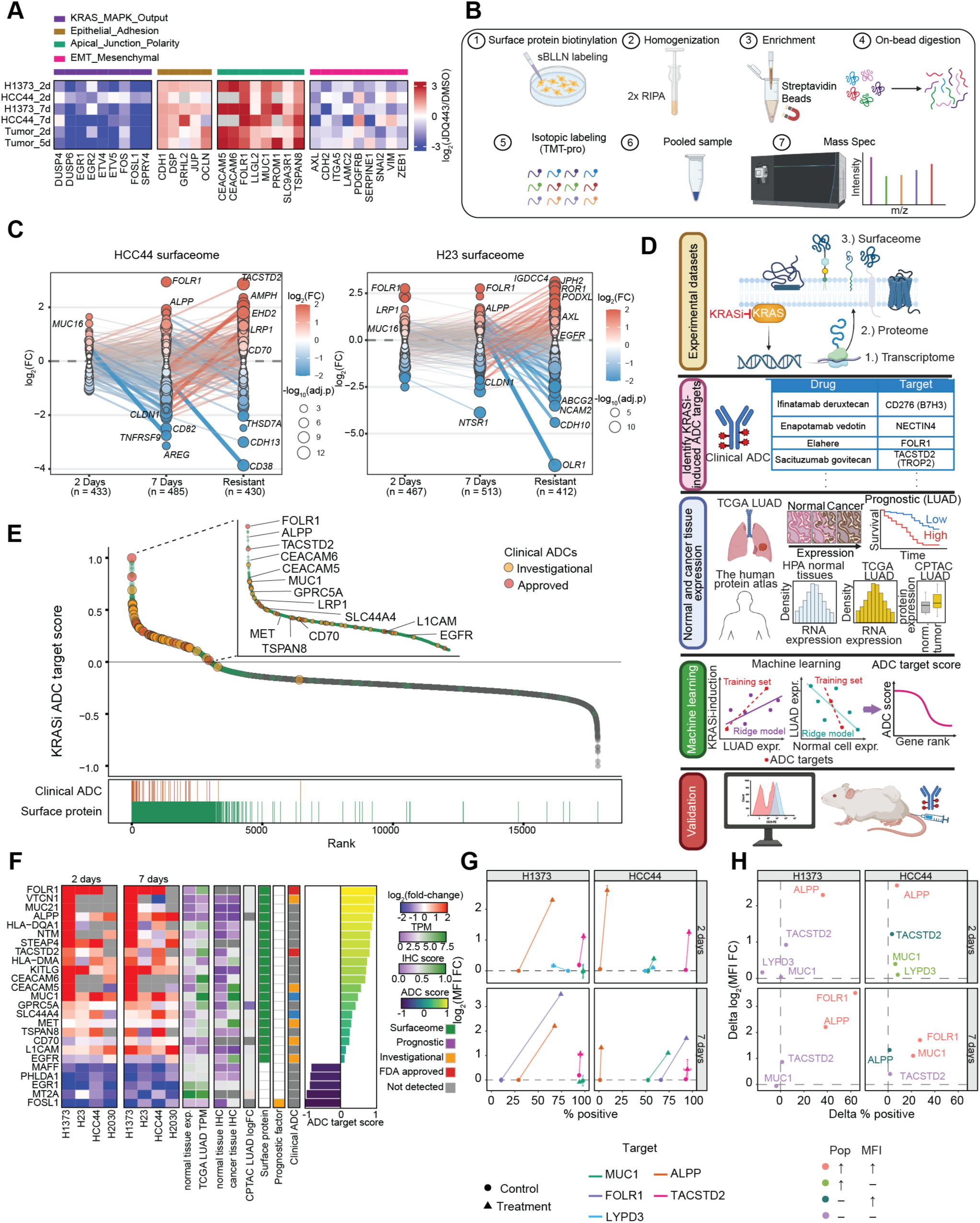
KRASi alters the NSCLC cell surfaceome with upregulation of ADC targets. **A.** Heatmap depicting expression profile for cell state related genes with KRASi treatment. **B.** Schematic for surfaceome enrichment using a biotinylation enrichment method. **C.** Trajectory plots illustrating differential expression of surface-annotated genes in HCC44 and H23 cells treated with JDQ443 for 2 days, 7 days and to the point of resistance. Plots display the log_2_(fold-change) of gene expression using integrated transcriptomic, proteomic, and surfaceome datasets using Stouffer’s weighted z-score. Points are sized by -log_10_(adjusted p value) of integrated data, and lines connecting genes between time points are sized and colored by the magnitude of change between the two points (D log_2_ fold-change). The number of significantly dysregulated (|log_2_(fold-change)| > 0.5, adjusted p value < 0.05) surface genes are shown below x-axis labels. **D.** Schematic for the workflow to identify and validate top surface-expressed targets by leveraging experimental datasets following KRASi, identifying the training set for machine learning using known clinical ADC targets, integrating normal and cancer cell expression datasets, training a ridge regression model to predict priority scores for ADC targets, and final validation. **E.** Rank plot of assigned KRASi ADC target score for all genes detected in transcriptomics and proteomics datasets. Surface proteins are annotated in green and proteins that have been clinically investigated or approved are orange or red, respectively. Barcode plots annotate the rank of corresponding surface proteins or clinically investigated ADC targets. Inset shows top KRASi ADC targets. **F.** Heatmap of select top ADC targets including known surface targets in lung cancer, and the bottom five scoring targets, showing the log_2_ fold-change for each target in each cell line for 2 day- or 7 day-treatment with JDQ443 using the integrated datasets. Features used to train the machine learning algorithm are shown, representing the normal and TCGA LUAD tissue RNA expression (log_2_(TPM +1), normal tissue and lung cancer IHC score from the HPA, log_2_ fold-change of tumor to normal adjacent tissue in lung cancer from CPTAC, surface protein annotation, significant prognostic indicator for survival in TCGA LUAD dataset. Clinical ADC designation and the KRASi ADC target score are shown (right), but were not included in training the ridge regression model. **G.** Arrow plot comparing MFI and percent positive cells for targets listed upon JDQ443 (100 nM) treatment for 2 and 7 days. Dots represent DMSO, triangles represent JDQ443 treated cells, arrows connect each cell line’s DMSO baseline to its JDQ443 endpoint. Error bars denote SD. **H.** Comparison of delta MFI and delta percent positive cells upon KRASi. Top right quadrant indicates a shift in both population and MFI and targets of interest.

We first compared our surface-enriched dataset to our global proteomics and transcriptomics datasets to determine the agreement and overlap. As expected, we found high concordance between our surface-enriched proteome and global proteome, demonstrating a high correlation across timepoints (Supplementary Fig. S6A, Supplementary Table S8). Annotation of proteins only detected by one modality showed that global proteomics detected more surface-annotated^41,42^ proteins, but nearly all dysregulated proteins were detected in both datasets. We then compared surface-annotated proteins detected by proteomics to those detected by transcriptomics. We found high agreement between modalities. Unsurprisingly, transcriptomics had greater coverage than global proteomics, detecting >2,000 surface proteins and identified more differentially regulated surface targets than global proteomics alone (Supplementary Fig. S6B).

The high agreement across platforms supports the robustness of our multi-omic approach. To maximize coverage of the KRAS-regulated surfaceome, we integrated our surface-enriched proteomics with global proteomic and transcriptomic datasets. We performed Stouffer’s weighted Z method to combine results from our three independent datasets to determine the most robust KRAS-regulated surfaceome detected in at least two modalities (Fig. 2C, Supplementary Fig. S6C, Supplementary Table S9). We found broad dysregulation of the cell surfaceome across time points and cell lines. The changes in the surfaceome were largely conserved at acute time points (i.e. 2 days and 7 days) but diverged substantially upon resistance. This suggests that distinct cell surface targets are presented upon initial KRAS targeting compared to the acquisition of therapeutic resistance. We annotated cell surface proteins with clinically investigated antibody drug conjugates^17^, and found several candidates that varied expression with temporal KRASi (Supplementary Fig. S6C).

To extend our evaluation of therapeutically tractable cell surface proteins beyond those previously targeted by ADCs, we developed a machine learning (ML) approach to systematically rank candidates (Figure 2D, Supplementary Table S10). We integrated public datasets of transcriptomic and protein expression data from the human protein atlas^43,44^ and the cancer genome atlas^45^ (TCGA) together with our robust multi-omics analysis across cell lines and xenografts treated with JDQ443. We trained a logistic ridge function using ten distinct features across all genes detected in at least two of our multi-omics datasets. These features included the transcriptional expression of a given gene in normal tissues^44^, immunohistochemistry (IHC) score in cancer tissues^44^, IHC score in normal tissues^44^, prognostic factor in lung adenocarcinoma patients^44^, transcript expression in TCGA^45^, and protein expression in lung adenocarcinoma samples from the clinical proteomic tumor analysis consortium (CPTAC)^46^. The model was trained on 32 clinically investigated ADC targets that we found upregulated in at least one of our multi-omics datasets following 2 days or 7 days KRASi. We predicted the log-odds probability of a given gene to be a tractable surface target to give a “KRASi ADC target score” using the coefficients of this model (Supplemental Fig. S6D). Evaluating all surface proteins by this ADC score revealed top ranked candidates including those that have been evaluated as ADC targets in lung cancer, such as FOLR1, ALPP, TACSTD2 (TROP2), and MUC1 (Fig. 2E, 2F).

We validated top candidates using flow cytometry in H1373 and HCC44 cells across treatment timepoints, enabling single cell resolution assessment of surface protein expression and upregulation. Confirmed targets included MUC1, ALPP, TROP2, and LYPD3 at 2 days and FOLR1, MUC1, ALPP, and TROP2 at 7 days (Fig. 2G). Notably, a subset of targets, including FOLR1 and MUC1 exhibited concordant increases in both the fraction of positive cells and mean fluorescence intensity (MFI) following KRASi, indicating a population wide shift in surface expression rather than selective expansion of a preexisting high expressing subpopulation (Fig. 2H). Based on the magnitude of its KRASi dependent upregulation, FOLR1 was prioritized for further functional and therapeutic characterization.

### FOLR1 expression is inversely associated with KRAS activity in lung adenocarcinoma

Given that FOLR1 emerged as a top candidate from our multi-omics analysis and was validated by flow cytometry, we next evaluated its baseline expression landscape in lung cancer^47^. Integration of TCGA transcriptomic and CPTAC proteomic data for patients with lung adenocarcinoma showed high correlation of protein abundance with FOLR1 RNA expression, suggestion RNA levels accurately model protein expression in patient samples (Supplementary Fig. S7A). We next assessed baseline FOLR1 expression in KRAS^G12C^ mutant tumors from the CodeBreaK 100/200 trials, comprising 234 patients with previously treated advanced NSCLC^48^ and an independent cohort of KRAS^G12C^ mutant NSCLC from the KRYSTAL-1 study (n = 68)^3^. We found broad expression of FOLR1 across patient samples in both cohorts, ranging from high expression to low or nearly undetectable (Supplementary Fig. S7B-C). To determine molecular pathways associated with FOLR1 expression, we classified patient samples as FOLR1-high or FOLR1-low and performed differential expression analysis and GSEA. We found strong depletion of KRAS/ERK-regulated genes in samples with high FOLR1 expression, and enrichment of KRAS/ERK-suppressed genes in FOLR1-low samples, suggesting the strongest determinant of FOLR1 expression is KRAS signaling output (Fig. 3A). This is consistent with our observation that FOLR1 expression is induced with KRASi. Additionally, we found enrichment of gene sets related to surfactant metabolism, consistent with an alveolar epithelial (AT2) identity, as well as mucin-associated glycosylation pathways, in patients with high FOLR1 expression (Supplementary Fig. S7D). Notably, these samples were derived from post-progression, heavily pretreated patients, indicating that FOLR1 expression is maintained in advanced, treatment-refractory KRAS^G12C^ NSCLC.

**Figure 3.**
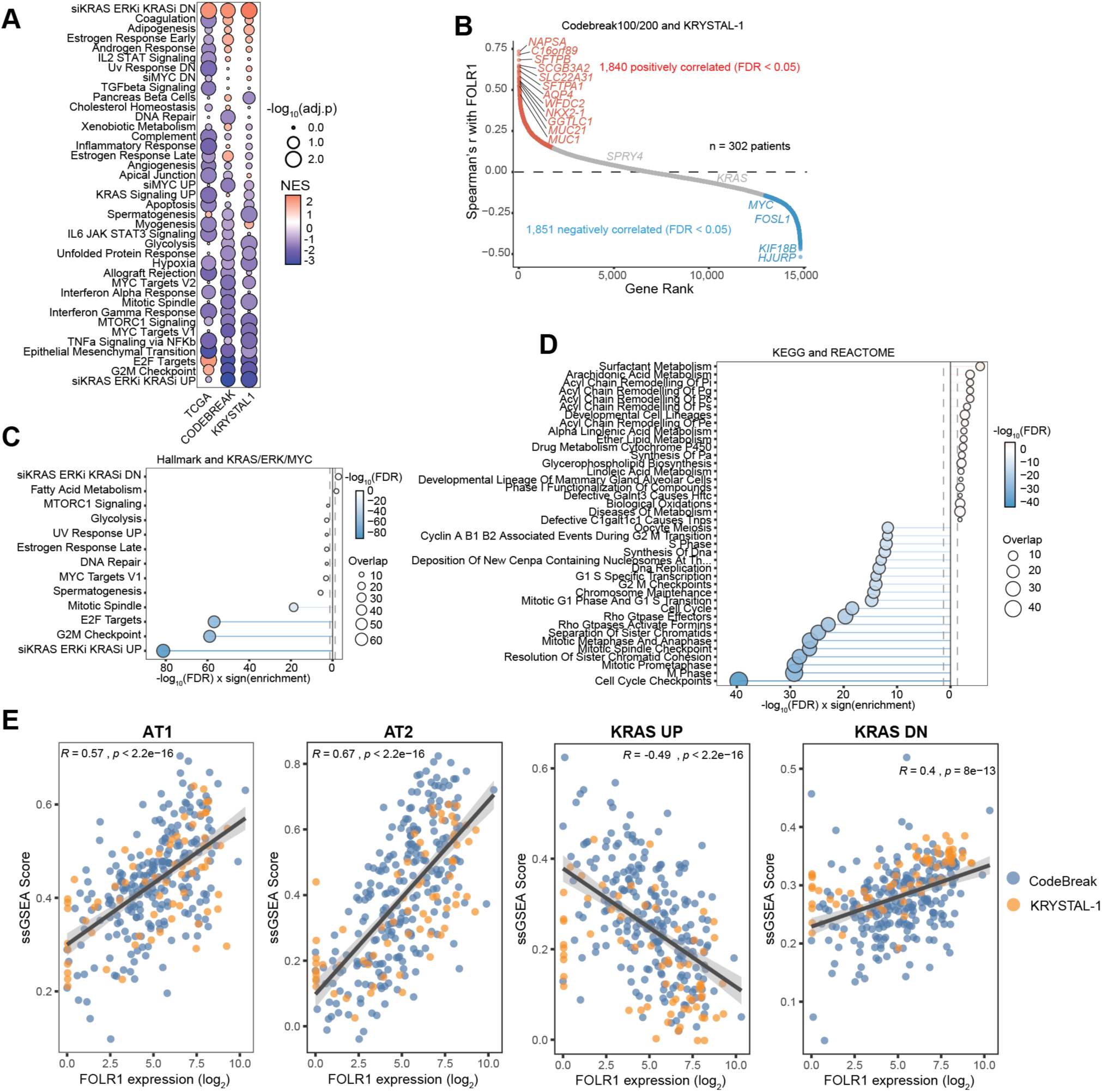
FOLR1 is detected in lung cancer and has inverse correlation with KRAS pathway activity. **A.** GSEA of differential gene expression between FOLR1-high and FOLR1-low patient samples from TCGA data of lung adenocarcinoma patients, and KRAS^G12C^-mutant NSCLC from CodeBreaK 100/200 trial or KRYSTAL-1 trial. Negative enrichment represents pathways depleted in samples with high FOLR1 expression and positive enrichment represents pathways increased in FOLR1-high samples compared to FOLR1-low. **B.** Rank plot showing spearman correlation coefficients of RNA expression with FOLR1 expression in KRAS^G12C^-mutant NSCLC patient samples from CodeBreaK100/200 and KRYSTAL-1 trials, with genes positively correlated with FOLR1 (FDR<0.05) in red and negatively correlated in blue. **C.** Overrepresentation analysis (ORA) of positive or negative correlated genes by Fisher’s exact test using Hallmark of KRAS/ERK- and MYC-regulated gene sets. All significant gene sets are shown (dotted line, FDR<0.05), plotted with the -log_10_ false discovery rate signed by the direction of enrichment, with negative corresponding to genes negatively correlated with FOLR1 expression and positive corresponding to genes correlated FOLR1. **D.** ORA as in (**C**) using KEGG and REACTOME gene sets. **E.** Correlations of single sample GSEA (ssGSEA) enrichment of AT1, AT2, or KRAS/ERK-regulated transcriptional signatures versus FOLR1 expression in patient samples from CodeBreaK100/200 (blue) or KRYSTAL-1 (orange) trials.

To identify genes that are co-expressed with FOLR1, we performed cross correlation analyses for all other genes with FOLR1 across all patients within the CodeBreaK and KRYSTAL-1 cohorts (Fig. 3B, Supplementary Fig. S7E). We found 1,840 and 1,851 genes positively or negatively correlated with FOLR1 expression, respectively. Consistent with our GSEA analysis, we found genes involved in metabolic pathways that support the production of surfactants and mucins correlated with FOLR1, and cell cycle regulatory genes such as MYC and FOSL1 were negatively correlated. Overrepresentation analysis (ORA) of the top positively and negatively correlated genes agreed with our GSEA results, showing KRAS/ERK-regulated genes are the most significant gene set negatively correlated with FOLR1 expression (Fig. 3C). This further demonstrates that KRAS activity is anticorrelated with FOLR1 expression and that KRASi is predicted to induce FOLR1, consistent with our proteomics data. Consistent with the surfactant pathway enrichment observed in FOLR1 high patient cohort (Supplementary Fig. S7D), we found that FOLR1 is co-expressed with genes encoding pulmonary surfactant proteins (SFTPA1/B/C/D) with an enrichment of lipid metabolic pathways supporting surfactant biosynthesis (Fig. 3D), together reflecting an AT2 differentiation program^49^. Additionally, FOLR1 expression was found to be highly correlated with the cell lineage factor NKX-2.1 (Fig. 3B), which dampens ERK/MAPK signaling via DUSP6 downstream of KRAS^50^ and drives both AT2 differentiation, the well-established cell of origin for KRAS-driven lung adenocarcinoma^51^, and AT1 differentiation^52^, a cell type recently shown to also serve this role^53^.

Our correlation analysis from KRAS^G12C^-mutant lung cancer patients showed FOLR1 expression is tightly connected to KRAS activity and the epithelial differentiation state of lung cancer. To more directly test the link to alveolar epithelial states, we computed single-sample GSEA (ssGSEA) scores for published AT1 and AT2 gene signatures^54^ for individual patient samples. We observed a strong positive correlation between FOLR1 expression and both AT1- and AT2-like signature scores, indicating that FOLR1 marks a lung epithelial cell state enriched for alveolar differentiation and surfactant metabolism (Fig. 3E). Extending this analysis to KRAS/ERK-regulated transcriptional programs showed a significant negative correlation with KRAS activity and positive correlation with KRAS pathway suppression (Fig. 3E). Collectively, these findings identify FOLR1 as a broadly expressed surface protein whose levels are inversely associated with oncogenic KRAS activity in lung cancer. This raises the possibility that KRAS signaling suppresses FOLR1 expression, and that KRASi may in turn upregulate FOLR1, potentially expanding patient eligibility for FOLR1-targeted therapies.

### Surface expression of FOLR1 is upregulated in response to RAS inhibitors

To systematically evaluate the impact of KRASi on FOLR1 surface expression, we selected a panel of KRAS^G12C^ mutant lung cancer cell lines with high, moderate, and low basal FOLR1 expression levels based on log_2_(TPM+1) expression data from DepMap (Supplementary Fig. S8A). After 2 days of JDQ443 treatment, there was a significant increase in MFI and percent positive cells in five out of the six lung cancer cell lines (Fig. 4A-C, Supplementary Fig. S8B).

**Figure 4.**
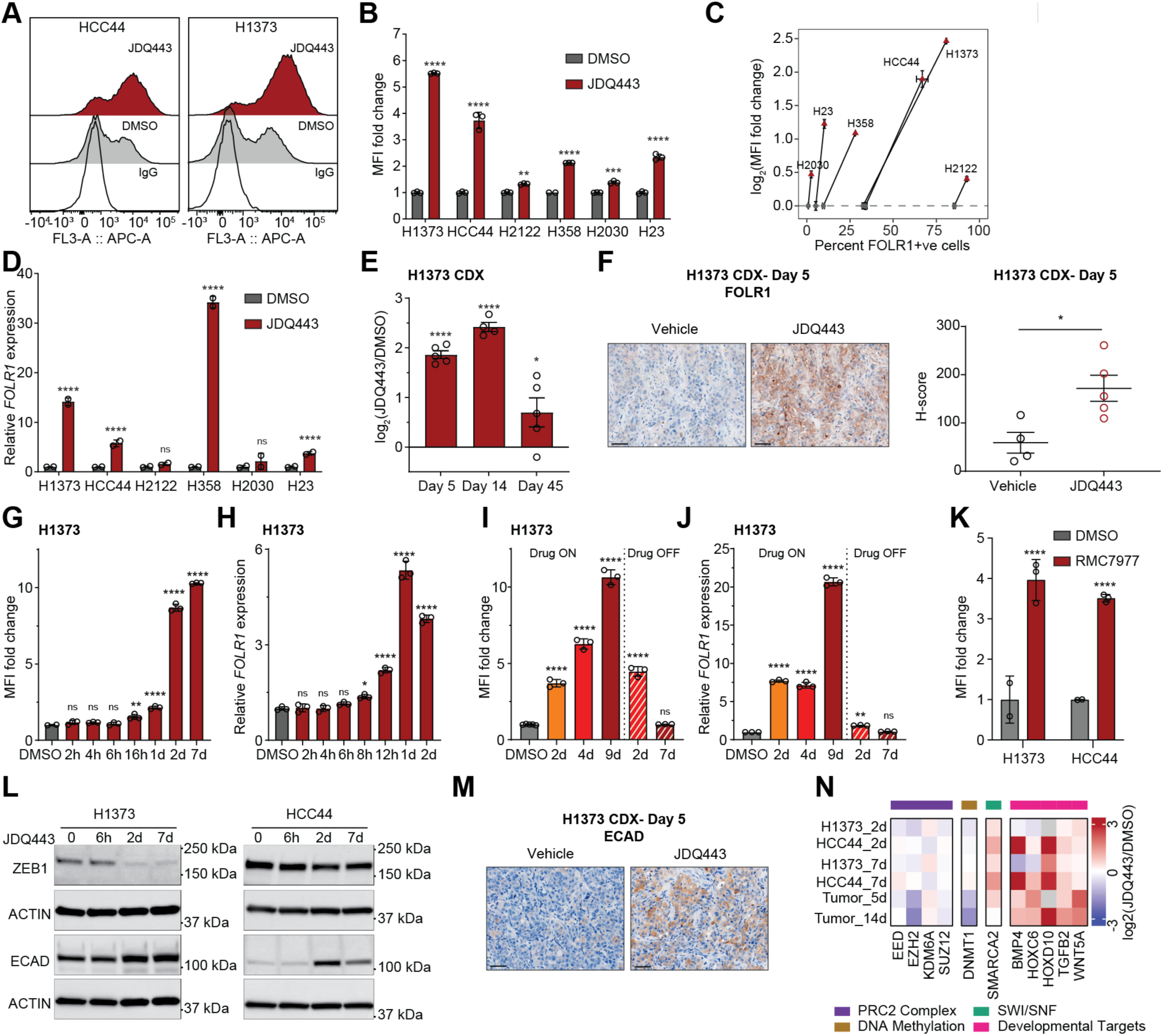
FOLR1 is upregulated in response to KRASi. **A.** Representative histograms for flow cytometry analysis of FOLR1 in H1373 and HCC44 cells treated with DMSO or JDQ443 (100 nM) for 2 days. Human IgG used as negative control. **B.** Mean fluorescence intensity (MFI) for FOLR1 in KRAS^G12C^ mutant lung cancer cell lines treated with DMSO or JDQ443 (100 nM) for 2 days. MFI was normalized to DMSO, data represented as mean ± SD. Statistical significance was determined by two-way ANOVA with Bonferroni’s multiple comparisons test. **C.** Arrow plot comparing MFI and percent positive cells for FOLR1 upon JDQ443 (100 nM) treatment for 2 days. Grey dots represent DMSO, red triangles represent JDQ443 treated cells, arrows connect each cell line’s DMSO baseline to its JDQ443 endpoint. Error bars denote ± SD. **D.** Relative FOLR1 expression determined by RT-PCR in KRAS^G12C^ mutant lung cancer cell lines treated with DMSO or JDQ443 (100 nM) for 2 days. Expression was normalized to DMSO, data represented as mean ± SD. Statistical significance was determined by two-way ANOVA with Bonferroni’s multiple comparisons test. **E.** Log_2_fold change in FOLR1 expression by RNA-seq in H1373 CDX tumors isolated at day 5, 14, and 45 post-JDQ443 (100 mg/kg) treatment, normalized to vehicle. **F.** *Left*: Immunohistochemistry for FOLR1 in H1373 tumors collected at day 5 post-treatment (JDQ443, 100 mg/kg). Scale bar: 50 µm. *Right*: H-score of FOLR1 expression determined by QuPath. Data represented as mean ± SEM. Statistical significance was determined by unpaired t-test. **G.** MFI fold change relative to DMSO over a time course in H1373 cells treated with JDQ443 (100 nM). Data represented as mean ± SD. Statistical significance was determined by ordinary one-way ANOVA. **H.** Relative FOLR1 expression determined by RT-PCR over a time course in H1373 cells treated with JDQ443 (100 nM) or DMSO. Expression was normalized to DMSO, data represented as mean ± SD. Statistical significance was determined by ordinary one-way ANOVA. **I.** Drug wash-off experiment. H1373 cells treated with JDQ443 (100 nM) for (i) 2, 4, or 9 days continuously, (ii) 7 days followed by 2 days without drug, or (iii) 7 days followed by 7 days without drug. FOLR1 expression determined by flow cytometry. Data represented as mean ± SD. Statistical significance was determined by ordinary one-way ANOVA. **J.** Wash-off experiment as (I), with FOLR1 expression determined by RT-PCR. Data represented as mean ± SD. Statistical significance was determined by ordinary one-way ANOVA. **K.** MFI fold change relative to DMSO for HCC44 and H1373 cell lines treated with RMC-7977 (10 nM). Data represented as mean ± SD. Statistical significance was determined by ordinary one-way ANOVA. **L.** Western blots for H1373 and HCC44 cells treated with JDQ443 (100 nM) over a time course, probed for indicated markers. **M.** Immunohistochemistry for E-cadherin in H1373 tumors collected at day 5 post-treatment. Scale bar: 50 µm. **N.** Heatmap depicting expression of genes associated with chromatin response. Significance for all panels: *p < 0.05, **p < 0.01, ***p < 0.001, ****p < 0.0001.

As FOLR1 was upregulated in our transcriptomic data at 2 days (Supplementary Fig. S8C), we confirmed upregulation of FOLR1 transcript with KRASi in our panel of KRAS^G12C^ mutant cell lines (Fig. 4D). Based on our transcriptomics data from H1373-CDX model, FOLR1 was upregulated on days 5, 14, and 45 of treatment with JDQ443 (Fig. 4E). We further confirmed FOLR1 upregulation by immunohistochemistry (Fig. 4F). In a time-course assay, we observed that FOLR1 surface expression increased between 16 and 24 hours following the start of KRASi treatment and was maintained at 7 days of treatment (Fig. 4G). Temporal profiling showed that FOLR1 transcript upregulation was detectable as early as 12 hours post-treatment (Fig. 4H).

To determine if FOLR1 expression was sustained after drug withdrawal, we treated H1373 and HCC44 cells with KRASi for 2 days followed by drug wash-off for 2 days and 7 days (Fig. 4I-J and S8D-E). FOLR1 transcript levels decreased upon drug withdrawal as measured at 2 days and 7 days whereas surface expression was maintained at 2 days off drug but decreased to baseline levels by 7 days in absence of drug (Fig. 4I-J, Supplementary Fig. S8D-E). These results indicate that FOLR1 expression upon KRASi treatment is transcriptionally regulated and continuous exposure to KRASi is required for sustained cell surface expression.

To further demonstrate broad applicability beyond mutant-specific KRASi, we evaluated the effect of the RAS(ON) multi-selective tricomplex inhibitor (RMC7977) which has previously been shown to be more effective in NSCLC^11^. In KRAS^G12C^ mutant lung cancer cell lines, there was a significant increase in FOLR1 expression in response to RMC7977 (Fig. 4K). Next, we tested FOLR1 expression in KRAS mutant ovarian cancer cells (OVCAR5 (KRAS^G12V^) and SNU8 (KRAS^G12D^)) with RMC7977 treatment (Supplementary Fig. S8F). There was a significant upregulation of FOLR1 in ovarian cancer cells RMC7977 both at surface and transcript levels (Supplementary Fig. S8G, S8H).

Having established FOLR1 upregulation as a consequence of (K)RAS inhibition in KRAS mutant lung and ovarian cancer, we confirmed this phenomenon in independent datasets and asked whether it extends beyond KRASi to other MAPK-pathway inhibitors and tumor contexts. Analysis of publicly available transcriptomic datasets spanning KRAS and EGFR inhibition across cell lines, patient-derived xenografts, and patient biopsies revealed frequent FOLR1 induction upon MAPK-pathway suppression, particularly in vitro and in PDX models (Supplementary Fig. S8I-J). FOLR1 induction was more variable in patient biopsies, increasing in some KRASi- and EGFRi-treated patients but decreasing in others. This heterogeneity may reflect factors not captured by the more controlled in vitro and PDX systems, including biopsy sampling after cessation of KRAS inhibition.

FOLR1 has been reported to promote activation of ERK1/2 signaling in multiple cancer contexts, including cervical cancer^55^ and colon cancer^56^. To determine if FOLR1 upregulation upon KRASi could be a component of resistance in NSCLC, we generated FOLR1 knockout cells using CRISPR/Cas9 genome editing (Supplementary Fig. S9A). Loss of FOLR1 did not alter sensitivity of lung cancer cells to KRASi (Supplementary Fig. S9B). These results suggest that while FOLR1 is upregulated following KRASi, it does not mediate resistance in this context.

Basal levels of folate in standard cell culture media (2.3 µM) are significantly higher than physiological levels (3-50 nM). Although FOLR1 is not a primary means of folate uptake in cells, excess folate in media can drive intracellular signaling pathways such as JAK-STAT3 and ERK1/2 that may not occur at physiological folate concentrations^21^. To ensure that FOLR1 upregulation with KRASi was not due to excess folate in the media, we cultured the cells in RPMI with no folate and dialyzed FBS and supplemented with increasing concentrations of folate. When treated with KRASi, there was consistent upregulation of FOLR1 in lung cancer cells with no significant difference between conditions with different folate levels in the media (Supplementary Fig. S9C).

FOLR1 has non-canonical functions beyond folate transport in development and epithelial morphogenesis^57–59^. FOLR1 localized on the apical surface of neural epithelial cells mediates folic acid uptake to promote apical constriction required for neural epithelial morphogenesis^59^. FOLR1 also balances apical trafficking and maintains cell-cell adhesion and apical-basal polarity by interacting with and impeding endocytosis of C-cadherin^57,58,60^. Overall, FOLR1 is closely linked to epithelial identity structurally and functionally. Therefore, FOLR1 upregulation upon KRASi likely reflects a coordinated shift toward an epithelial gene expression program. We confirmed an induction of epithelial transition at the protein level by increased E-cadherin and decreased ZEB1 expression, starting at 2 days and sustained through 7 days of KRASi treatment (Fig. 4L) which was confirmed in H1373 CDX after 5 days of JDQ443 treatment (Fig. 4M).

Chromatin dynamics are important regulators of epithelial cell identity and cellular plasticity^61^. Previous studies have shown that RAS signaling can engage epigenetic mechanisms to silence genes either by hypermethylation of CpG islands, or recruitment of DNMT1 or the PRC2 complexes to gene promoters^62–65^. Oncogenic KRAS signaling has been shown to suppress epithelial differentiation and identity in multiple contexts^10,66–68^. Therefore, KRASi could restore a more differentiated, epithelial phenotype as a part of chromatin derepression. Consistent with this, KRASi was associated with broad attenuation of chromatin repression across multiple regulatory layers. Core components of the PRC2 complex (EZH2, SUZ12, EED) were reduced, alongside diminished DNA methylation maintenance (DNMT1), suggesting a shift toward a more accessible chromatin landscape. In parallel, increased expression of the H3K27 demethylase KDM6A and the chromatin remodeling factor SMARCA2 further support active chromatin opening (Fig. 4N). Repressive chromatin mechanisms have been implicated in silencing epithelial identity genes such as E-cadherin^69–72^. The reduction in PRC2 components and DNMT1 following KRASi is consistent with relief of these repressive mechanisms. We also observed upregulation of developmentally silenced target genes, including HOX family members (HOXC6, HOXD10) and key signaling mediators (WNT5A, BMP4, TGFB2), suggesting that KRASi drives broad epigenetic reprogramming. Since FOLR1 has an established structural and functional role related to epithelial polarity and adhesion, the coordinated upregulation of FOLR1 and E-cadherin is consistent with an epithelial state shift upon KRASi.

### KRASi enhances the cellular uptake and antitumor activity of the FOLR1-ADC

Having validated FOLR1 upregulation as a conserved response to KRAS/MAPK pathway inhibition, we next evaluated the effects of targeting FOLR1 using antibody drug conjugates with and without KRASi. We first tested the baseline sensitivity of lung and ovarian cancer cell lines to mirvetuximab soravtansine-gynx (Fig. 5A). Sensitivity to mirvetuximab soravtansine-gynx directly correlated with baseline antibody binding capacity (Fig. 5B-C) and this was not due to differential sensitivity to DM4, the cytotoxic payload of mirvetuximab soravtansine-gynx (Supplementary Fig. S10A-B).

**Figure 5.**
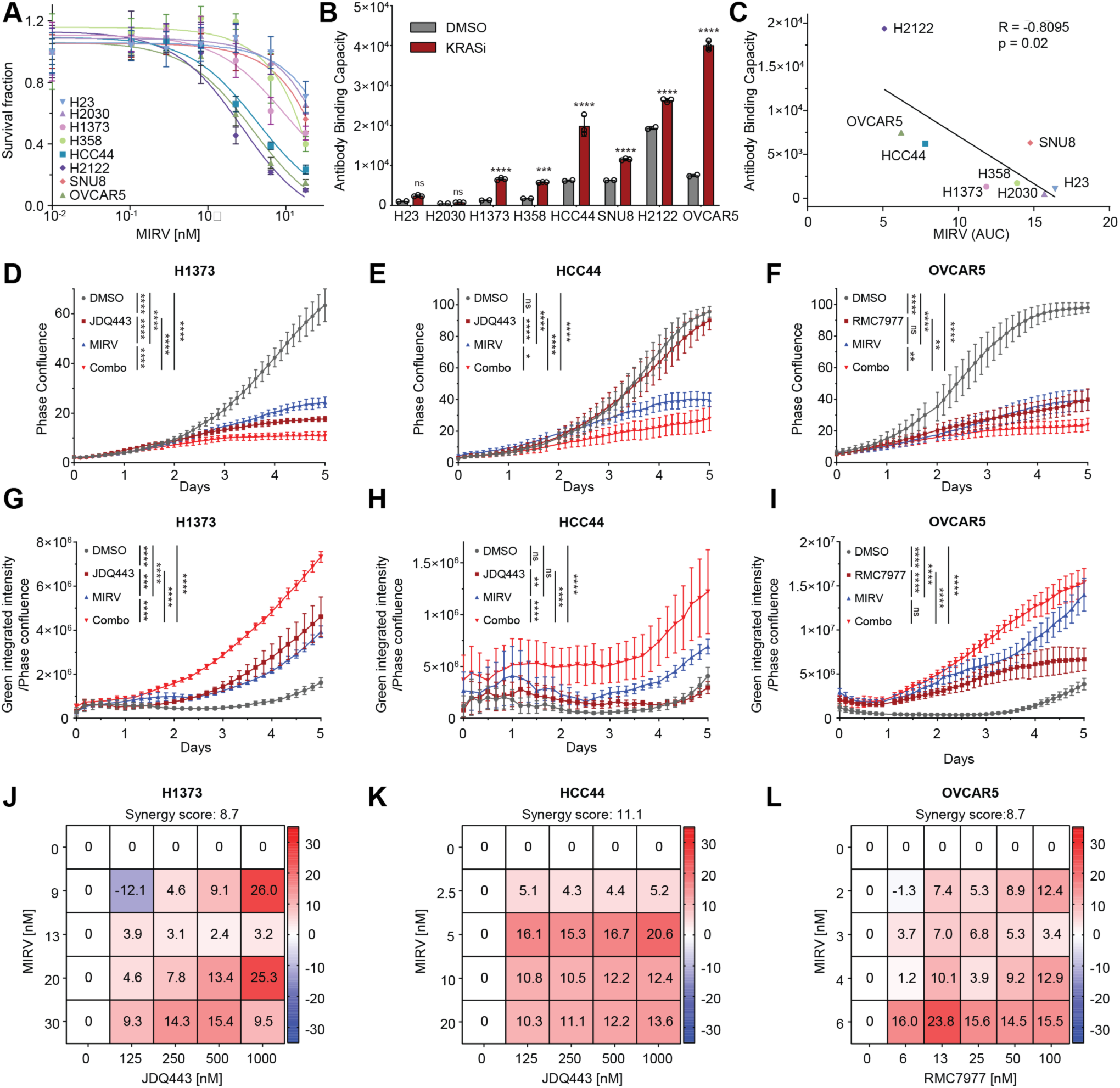
Combination KRASi and FOLR1-ADC demonstrates efficacy in NSCLC cell culture models. **A.** Dose response curves for KRAS mutant lung and ovarian cancer cells for mirvetuximab soravtansine-gynx (MIRV) for 2 days. Cell viability was determined by CellTiter-Glo assay and normalized to DMSO. Data represented as mean ± SD. **B.** Antibody binding capacity (ABC) assay for KRAS mutant lung and ovarian cancer cells for FOLR1 at 2 days. Data represented as mean ± SD. Statistical significance was determined by two-way ANOVA with Bonferroni’s multiple comparison tests. **C.** Spearman correlation of ABC and area under the curve (AUC) for MIRV (calculated in GraphPad PRISM). **D-F.** Incucyte based growth assay for H1373 (left), HCC44 (center) and OVCAR5 (right) cells treated DMSO, JDQ443 (100 nM), MIRV (5 nM), or combination over 5 days. Data represented as mean ± SD. Statistical significance was determined by two-way ANOVA. **G-I.** Cell death determined by Incucyte Cytotox Dye for Counting Dead Cells (green) for H1373 (left), HCC44 (center) and OVCAR5 (right) cells treated with DMSO, JDQ443 (100 nM), MIRV (5 nM), or combination for 5 days. Data represented as mean ± SD. Statistical significance was determined by two-way ANOVA. **J-L.** Synergy assay for H1373 (left), HCC44 (center), and OVCAR5 (right) cells across a range of doses. Synergy score calculated by the HSA (highest single agent) method. Significance for all panels: *p < 0.05, **p < 0.01, ***p < 0.001, ****p < 0.0001.

Quantitative analysis of FOLR1 binding to cells upon KRASi revealed an increased number of surface FOLR1 molecules across different cell lines (Fig. 5B). Additionally, an increased internalization of mirvetuximab antibody was observed in lung and ovarian cancer cells treated with KRASi without any effect on IgG internalization (Supplementary Fig. S10C). In growth assays over 5 days, the combination of KRASi and mirvetuximab soravtansine-gynx showed greater efficacy in H1373, HCC44 and OVCAR5 cells (Fig. 5D-F). Cells treated with the combination also showed greater cell death than with single agents at day 5 (Fig. 5G-I). Furthermore, KRASi and mirvetuximab soravtansine-gynx combination was synergistic across a range of doses in H1373, HCC44 and OVCAR5 cells (Fig. 5J-L, Supplementary Fig. S10D). As DM4 is a microtubule inhibitor, combination treatment of KRASi and mirvetuximab soravtansine-gynx led to an increase in pH2AX accumulation over 5 days indicative of a DNA damage response (Supplementary Fig. S10E). We next tested the combination efficacy of KRASi and mirvetuximab soravtansine-gynx in cells with very low basal levels of FOLR1 and minimal increase in KRASi-induced FOLR1 expression (H23 and H2030). There was no combination efficacy (Supplementary Fig. S10F-G) highlighting the importance of basal FOLR1 expression for a KRASi and FOLR1-targeting ADC therapeutic approach.

### Combination of KRASi and FOLR1-ADC provides sustained tumor suppression

We next evaluated the efficacy of combination KRASi and FOLR1-targeting ADC in multiple in vivo NSCLC models. Mice bearing H1373 xenografts were treated for 30 days and then monitored off treatment for tumor regrowth. JDQ443 alone led to significant tumor growth control on treatment consistent with previously published studies^73^. Mirvetuximab soravtansine-gynx monotherapy did not have a significant tumor growth control compared to vehicle arm, which was expected as H1373 have relatively low basal FOLR1 levels. Since FOLR1 expression was upregulated after 5 days of KRASi, we next sought to evaluate if mirvetuximab soravtansine-gynx combination efficacy would be different between concomitant or delayed mirvetuximab soravtansine-gynx treatment. Overall, combination of JDQ443 and mirvetuximab soravtansine-gynx was more effective than the monotherapies. However, there was no difference in tumor growth rates between the groups where mirvetuximab soravtansine-gynx was administered either concomitantly or delayed by 5 days (Fig. 6A, Supplementary Fig. S11A). Upon drug withdrawal, JDQ443 treated tumors regrew to baseline in ∼3 weeks whereas combination arm tumors reached a similar size in ∼6 weeks. Although H1373 tumors showed sensitivity to JDQ443 alone, we observed a deeper response in both combination arms at the end of treatment (Fig. 6B). Tumor doubling time for mice was greater in combination arms compared to monotherapies (Fig. 6C) and overall survival for mice treated with the combination was longer (Supplementary Fig. S11B).

**Figure 6.**
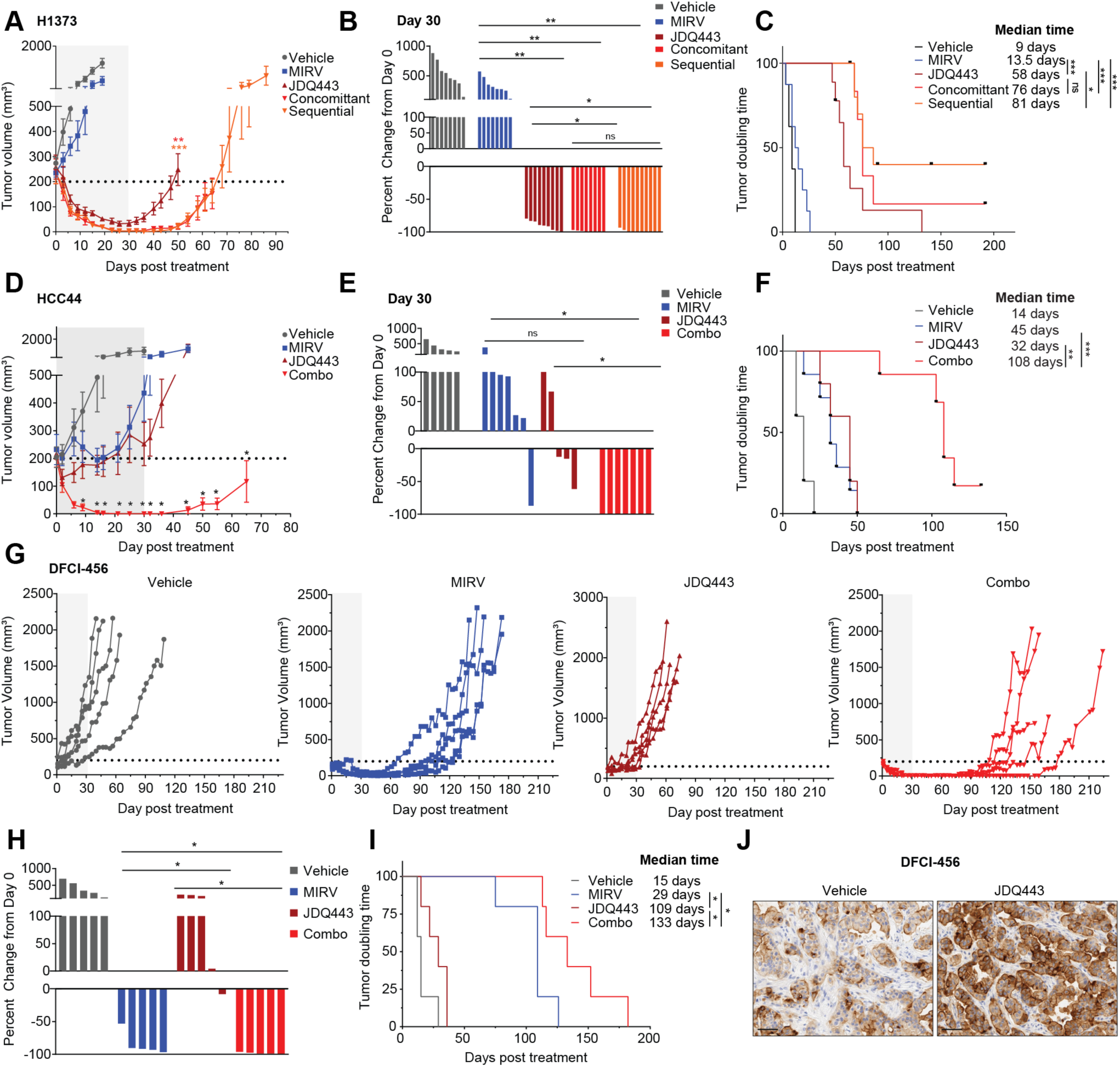
Combination KRASi and mirvetuximab soravtansine-gynx demonstrates efficacy in NSCLC tumor models. **A.** Tumor growth curves for mice bearing H1373 xenografts (n = 8-10 per arm) treated with vehicle, mirvetuximab soravtansine-gynx (MIRV, 5 mg/kg), JDQ443 (100 mg/kg), concomitant (JDQ443 and MIRV initiated on day 0), or sequential (MIRV initiated 5 days after JDQ443 treatment start). Data are represented as mean tumor volume ± SEM; curves are truncated at the timepoint when the first animal in each respective group was euthanized. Statistical significance among JDQ443, concomitant, and sequential arms at day 50 was determined by one-way ANOVA; vehicle and MIRV monotherapy arms were excluded from this comparison as tumors in these groups did not reach day 50. Grey box indicates the dosing period and the dashed horizontal line marks the average tumor volume (200 mm³) at treatment start. **B.** Waterfall plot depicting percent change in tumor volume for individual H1373 xenograft bearing mice treated with vehicle (n = 8), MIRV (5 mg/kg; n = 8), JDQ443 (100 mg/kg; n = 9), concomitant combination (JDQ443 and MIRV initiated on day 0; n = 8), or sequential combination (MIRV initiated 5 days after JDQ443 treatment start; n = 10). Response was evaluated at end of treatment (day 30), or at the end point for mice that did not reach day 30. Each bar represents one animal. Statistical significance was determined by Kruskal-Wallis test (p = 6.9 × 10^-7^) with pairwise Wilcoxon rank-sum tests and Holm correction. **C.** Kaplan-Meier curves showing time to tumor volume doubling relative to day 0 for mice bearing H1373 xenografts treated with vehicle (n = 8), MIRV (5 mg/kg; n = 8), JDQ443 (100 mg/kg; n = 9), concomitant combination (JDQ443 and MIRV initiated on day 0; n = 8), or sequential combination (MIRV initiated 5 days after JDQ443 treatment start; n = 10). Median time to doubling was 9, 13.5, 58, 76, and 81 days for vehicle, MIRV, JDQ443, concomitant, and sequential arms, respectively. Statistical significance was determined by log-rank test (overall p < 0.0001) with Holm-adjusted pairwise comparisons. **D.** Tumor growth curves for mice bearing HCC44 xenografts (n = 5-7 per arm) treated with vehicle, MIRV (5 mg/kg), JDQ443 (100 mg/kg) or combination. Data are represented as mean tumor volume ± SEM; curves are truncated at the timepoint when the first animal in each respective group was euthanized. Statistical significance between JDQ443 and combination arms was determined by one-way ANOVA as indicated on the graph. **E.** Waterfall plot depicting percent change in tumor volume for individual HCC44 xenograft-bearing mice treated with vehicle, MIRV (5 mg/kg), JDQ443 (100 mg/kg) or combination. Response was evaluated at end of treatment (day 30). Each bar represents one animal. Statistical significance was determined by Kruskal-Wallis test (p = 2.7 × 10^-4^) with pairwise Wilcoxon rank-sum tests and Holm correction. **F.** Kaplan-Meier curves showing time to tumor volume doubling relative to day 0 for mice bearing HCC44 xenografts treated with vehicle (n = 5), MIRV (5 mg/kg; n = 7), JDQ443 (100 mg/kg; n = 5) or combination (n = 7). Median time to doubling was 14, 32, 45 and 108 days for vehicle, MIRV, JDQ443, combination arms, respectively. Statistical significance was determined by log-rank test (overall p < 0.0001) with Holm-adjusted pairwise comparisons. **G.** Individual tumor volume trajectories over time for mice bearing DFCI-456 patient derived xenografts (PDX) treated with vehicle (n = 5), MIRV (5 mg/kg, n = 5), JDQ443 (100 mg/kg, n = 5), or combination (n = 5), shown as separate panels per arm. Each line represents an individual animal. Grey boxes indicate the dosing period, and the dashed horizontal line marks the average tumor volume (200 mm³) at treatment start. **H.** Waterfall plot depicting percent change in tumor volume for individual DFCI-456 PDX bearing mice treated with vehicle, MIRV (5 mg/kg), JDQ443 (100 mg/kg) or combination. Response was evaluated at end of treatment (day 30). Each bar represents one animal. Statistical significance was determined by Kruskal-Wallis test (Kruskal-Wallis p = 7.7 × 10^-4^) with pairwise Wilcoxon rank-sum tests and Holm correction. **I.** Kaplan-Meier curves showing tumor volume doubling relative to day 0 for mice bearing DFCI-456 PDX treated with vehicle (n = 5), MIRV (5 mg/kg; n = 5), JDQ443 (100 mg/kg; n = 5) or combination (n = 5). Median time to doubling was 15, 109, 29 and 133 days for vehicle, MIRV, JDQ443 and combination arms, respectively. Statistical significance was determined by log-rank test (overall p < 0.0001) with Holm-adjusted pairwise comparisons. **J.** Immunohistochemistry for FOLR1 on DFCI-456 PDX treated with JDQ443 for 5 days. Scale bar: 50 µm. **K.** Immunohistochemistry for Ki67 and yH2AX on DFCI-456 PDX treated with vehicle, JDQ443, MIRV and combination for 5 days. Statistical significance was determined by one-way ANOVA. Scale bar: 50 µm. Significance for all panels: *p < 0.05, **p < 0.01, ***p < 0.001, ****p < 0.0001.

Next, we evaluated a HCC44 xenograft model for combination efficacy. HCC44 tumors demonstrated a partial response to JDQ443 or mirvetuximab soravtansine-gynx monotherapy characterized by initial tumor stasis. In both monotherapy arms, tumor regrowth occurred in some mice during treatment and in all mice immediately following withdrawal of drug (Fig. 6D, Supplementary Fig. S11C). Remarkably, tumors treated with combination therapy demonstrated complete responses at the end of dosing (Fig. 6E) with regrowth delayed by ∼6 weeks compared to monotherapy treatments. Tumor doubling time for mice was significantly longer in the combination arm compared to monotherapies (Fig. 6F) and overall survival for mice treated with combination was significantly higher than either monotherapy (Supplementary Fig. S11D).

Next, we evaluated combination efficacy in a KRAS^G12C^ mutant PDX model with high basal FOLR1 expression. Mice were treated for 30 days and then monitored off treatment. Mirvetuximab soravtansine-gynx treatment alone had a significant effect on these tumors due to higher basal FOLR1 expression whereas these tumors were a priori resistant to KRASi. Combination therapy demonstrated a more rapid, deeper and sustained response. Furthermore, tumor relapse in the combination arm was delayed by ∼8 weeks compared to mirvetuximab soravtansine-gynx monotherapy arm (Fig. 6G). At the end of treatment, tumors in the combination arm had a greater tumor growth inhibition compared to mirvetuximab soravtansine-gynx monotherapy (Fig. 6H, Supplementary Fig. S11E). Tumor doubling time for mice in combination arm was significantly longer than the mirvetuximab soravtansine-gynx or JDQ443 monotherapy arms (Fig. 6I) whereas overall survival in mice treated with combination was not significantly longer than mirvetuximab soravtansine-gynx, likely due to increased tumor ulceration in the combination arm (Supplementary Fig. S11F). Combination of JDQ443 and mirvetuximab soravtansine-gynx was well tolerated in all mouse models as monitored by mouse weight (Supplementary Fig. S10G).

In a 5-day short-term pharmacodynamic study in the DFCI-456 PDX model, there was marked upregulation of FOLR1 in the JDQ443 treatment arm (Fig. 6J). Additionally, at day 5, there was a decrease in proliferating cells in the combination arm as determined by Ki67 staining and increased accumulation of yH2AX, indicating DNA damage in tumor cells (Fig. 6K). Combined, these results indicate that KRASi induces FOLR1 upregulation in vivo and upfront treatment of KRASi with a FOLR1-targeting ADC induces deep and sustained tumor regressions compared to either monotherapy.

## DISCUSSION

In this study, we used an integrated multi-omic approach to systematically characterize the adaptive landscape in response to KRAS^G12C^ inhibitors (KRASi) at early, late and resistance timepoints. Beyond identifying signaling adaptations, we also highlight and characterize a previously underappreciated link between KRASi and dynamic remodeling of the cell surface proteome. In particular, we identify folate receptor alpha (FOLR1) as a high-priority antibody drug conjugate (ADC) target upregulated upon KRASi and demonstrate combination efficacy of KRASi with a FOLR1-ADC.

KRASi have transformed the treatment of KRAS mutant lung cancer, with sotorasib and adagrasib demonstrating meaningful clinical benefit and now being evaluated in the first line setting [KRYSTAL-7, NCT06582771, KROCUS (NCT05756153)]. However, resistance to KRASi is nearly universal. Combinations targeting upstream regulators (SHP2, SOS1), downstream effectors (MEK, ERK), and cooccurring alterations have been pursued, yet durable disease control remains elusive and toxicity limits vertical pathway inhibition strategies^74–76^ Acquired resistance arises through alterations in both KRAS oncoprotein and alternate survival nodes, and intratumoral heterogeneity further limits the impact of targeting any single genetic resistance event^77,78^. This lack of convergent resistance mechanism underscores the importance of a deeper understanding of the non-genetic adaptive landscape.

Our proteomic, phosphoproteomic and transcriptomic evaluations across KRAS^G12C^ mutant models focused on tumor cell-intrinsic responses to KRASi treatment. Consistent with reduced proliferation, multi-omic profiling revealed a concurrent activation of stress and metabolic survival programs. Over time, these survival adaptations enabled cells to persist under continued KRASi pressure, providing a permissive context in which resistance-conferring alterations, such as secondary KRAS mutations, could emerge or expand. The resulting cell state was characterized by restoration of proliferation, specifically through reactivation of cyclin-dependent kinases (CDKs). Combination of CDK4/6 inhibitors have been evaluated in combination with KRASi, but resistance is common, further underscoring the limitations of targeting individual signaling nodes in isolation^73,79^.

A notable and underappreciated consequence of KRASi is the broad remodeling of tumor cell identity. KRAS-mutant lung cancers frequently occupy a relatively dedifferentiated and mesenchymal-like state maintained by active RAS–MAPK signaling, which suppresses epithelial and lineage-associated differentiation programs^10^. Pharmacologic KRASi transiently relieves these constraints, enabling re-expression of epithelial transcriptional programs and promoting an early shift toward an epithelial and apical cell state, consistent with our GSEA analyses and recent studies describing KRASi induced cellular plasticity and alveolar differentiation programs^8,10,11^. This cell state transition is accompanied by broad remodeling of the cell surface proteome, with upregulation of epithelial-associated surface proteins including CDH1, KRT8, OCLN, CLDN18 and MUC1. The KRASi induced surfaceome and epithelial phenotype therefore represents a therapeutically exploitable vulnerability. Using orthogonal approaches, we identified top therapeutic targets for combination with KRASi. These included FOLR1, TROP2, ALPP and MUC1. FOLR1 emerged as a high-priority ADC target for lung cancer patients due to its potential for rapid clinical translatability.

FOLR1 expression is generally restricted to epithelial tissues in normal physiology, where it localizes to the apical surface and physically interacts with components of adherens junctions including C-cadherin and β-catenin^21,57^. FOLR1 is predominantly overexpressed in tumors of epithelial origin^23^. In lung cancer, FOLR1 expression has been reported in the majority of lung cancer cases, with higher levels observed in lower histologic grades^80^, where tumors retain a more differentiated and epithelial like state^47,81^. Additionally, our analysis of patient data from the CodeBreak and KRYSTAL trials revealed an inverse correlation between KRAS pathway activity and FOLR1 expression. Our results indicate that combination of KRASi with a FOLR1-ADC provides sustained tumor growth control. In patient-derived xenograft (PDX) model, combination therapy resulted in earlier tumor regression compared to KRASi monotherapy. This provides a rationale for incorporating FOLR1-ADC combinations into concurrent treatment frameworks, especially since KRAS^G12C^ inhibitors are being considered in the first line setting [KRYSTAL-7 (NCT04613596), NCT06582771, KROCUS (NCT05756153)]. While combination of KRASi with chemotherapy is under active investigation, tolerability may be limited. Targeted delivery of cytotoxic drugs via ADCs could circumvent toxicity issues while maintaining potency. However, tolerability to ADCs can similarly be limited by toxicity and one limitation of our study is that murine FOLR1 is not recognized by mirvetuximab limiting the ability to fully model potential combination toxicities. Of note, nearly all mice receiving combination therapy underwent tumor recurrence after treatment was stopped indicating that a longer therapy course would be necessary in patients. However, all models in this study were immunocompromised and it is possible that in the setting of an intact immune system, residual tumor cells may have been cleared following the initial deep tumor regressions observed^82,83^. Furthermore, there is likely an opportunity to leverage triple combination of KRASi, FOLR1-ADC and immune checkpoint inhibition (ICI) to improve durability of response^84,85^.

Patient stratification is critical for efficacy of FOLR1-ADC in combination with KRASi. Our results indicate that baseline expression of FOLR1 is required for further upregulation upon KRASi, indicating that patient selection strategies will need to account for initial expression levels to maximize therapeutic benefit. Similarly, a recent study with RAS(ON) multi-selective inhibitor in pancreatic cancer showed an upregulation of HER2 which led to combination efficacy of RASi with HER2-ADC in three out of four models. No meaningful combination effect was observed in model that had very low initial HER2 expression^34^. However, patients with low but detectable FOLR1 may still be eligible, as next-generation FOLR1-ADCs with extended half-lives, bystander killing activity, and improved payloads lower the expression threshold required for efficacy^29,86,87^. Our data also suggests that continuous KRASi is critical to sustain FOLR1 surface expression, which has important implications for treatment scheduling. Given that ADCs are typically administered on a 3-week cycle, optimization of dosing intervals and sequencing will be necessary to balance efficacy and toxicity. Wash-off experiments and timing studies further support the existence of a therapeutic window in which KRASi primes tumor cells for enhanced ADC sensitivity. Our data extend these observations by showing that FOLR1 is not only a static biomarker but also a dynamically regulated target responsive to KRASi.

Beyond KRAS^G12C^ lung cancer, targeting FOLR1 upon KRASi may have broader applicability across KRAS-driven tumor types, and even extend to other MAPK driven tumors such as EGFR-mutant lung cancers^47^, colorectal cancer^88^, ovarian cancer^89^ and melanoma^90^ patients. Beyond FOLR1, our surfaceome dataset nominates additional KRASi-induced targets, including ALPP, MUC1^91^, and TROP2, that could be exploited through alternative ADCs, bispecific antibodies co-targeting FOLR1 and MUC1^92^ or CAR-T approaches directed at the KRASi-induced surface state, offering combinatorial strategies for patients with low or absent FOLR1 expression.

Finally, our surfaceome dataset nominates therapeutically relevant targets at the point of acquired resistance for future exploration including L1CAM, LRP1, and PODXL providing options for patients who have progressed on KRASi. This highlights the significance of temporal surfaceome profiling for matching patients to surface-directed therapies at each stage of disease evolution under KRASi therapy. In summary, our work identifies a mechanistically informed combination strategy in which KRASi induces a targetable cell surface state, enabling enhanced efficacy of FOLR1-directed therapies. These findings support further clinical investigation of KRASi and FOLR1-ADC combinations and optimization of combination regimens to improve outcomes in lung cancer.

## METHODS AND MATERIALS

### Cell culture

Human lung cancer cell lines were obtained from ATCC. All cell lines were maintained in a centralized cell bank, authenticated by assessment of cell morphology as well as short tandem repeat fingerprinting, and routinely inspected for Mycoplasma contamination using PCR. After thawing, cell lines were cultured for no longer than 30 days. Cell lines were maintained at 37°C with 5% CO_2_ and grown in RPMI 1640 supplemented with 10% FBS and 1% penicillin/streptomycin, unless otherwise specified.

### Generation of isogenic models of acquired resistance to JDQ443

JDQ443 resistant cell lines were generated by continuous dose escalation from 1 nM to 5 μML. The point of resistance was defined as the ability of cells to maintain stable growth rates comparable to DMSO-treated controls in the presence of JDQ443. Once resistance to JDQ443 was confirmed, resistant cell lines were routinely maintained in medium with 1 µM JDQ443.

### Drug sensitivity assays

Cells were seeded in 96-well plates (1000 cells/well) in complete media on day 0 and allowed to adhere overnight. The next day, cells were treated with indicated doses of JDQ443 (Novartis), RMC7977 (MedChemExpress), mirvetuximab soravtansine-gynx (DFCI Pharmacy), DM4 (MedChemExpress) or DMSO (Day 1). Cell viability was assessed after 2-5 days post treatment using CellTiter-Glo reagent (Promega, G9243) as indicated. Luminescence was measured using a CLARIOstar microplate reader (BMG LABTECH). Luminescence values were normalized to DMSO-treated controls. Dose inhibition curves were generated using three-parameter nonlinear regression in GraphPad Prism. Technical triplicates were conducted for each biological replicate, and a total of at least three biological replicates were done for each cell line. Area under the curve (AUC) was calculated according to GraphPad PRISM guidelines.

### Drug combination synergy assays

Cells were seeded in 96-well plates (1000 cells/well) in complete media on day 0 and allowed to attach overnight. The next day, cells were treated with JDQ443, RMC7977 or mirvetuximab soravtansine-gynx at indicated doses and combination. Cell viability was measured on day 5 with CellTiter-Glo reagent (Promega, G9243) using manufacturer’s instructions. Luminescence was measured using CLARIOstar (BMG LABTECH). Synergy analysis was performed using the highest single agent (HSA) model in SynergyFinder (https://synergyfinder.fimm.fi/). Mean HSA scores for each drug combination were used to generate synergy heatmaps in GraphPad PRISM.

### Growth curves

Cells were seeded in 96-well plates (1,000 cells/well) in complete media on day 0 and allowed to attach in incubator overnight. The following day, cells were treated with JDQ443, RMC7977 or mirvetuximab soravtansine-gynx at indicated concentrations. Cells were incubated for 30 minutes in incubator to equilibrate before imaging. Imaging and analysis were performed using IncuCyte Zoom/S3 imagers (Essen Biosciences) and plotted on GraphPad PRISM.

### Cytotoxicity assay

Cells were seeded in 96-well plates (1,000 cells/well) in complete media on day 0 and allowed to attach in incubator overnight. The following day, cells were treated with JDQ443, RMC7977 or mirvetuximab soravtansine-gynx at indicated concentrations prepared in media containing Incucyte Cytotox Dye at a final concentration of 250 nM. Imaging and analysis were performed using IncuCyte Zoom/S3 imagers (Essen Biosciences) and plotted on GraphPad PRISM.

### RNA isolation and quantitative real-time PCR

Cells were seeded in 6-well plates and treated with DMSO (control) or JDQ443 (100 nM) for the indicated time points. Cell pellets were collected for RNA extraction using TRIzol reagent (ThermoFisher, 15596026) following the manufacturer’s protocol. cDNA was synthesized using the High-Capacity cDNA Reverse Transcription Kit (Applied Biosystems, 4368814). qPCR was performed using SYBR™ Green Universal Master Mix (Applied Biosystems, 4309155) on an Applied Biosystems QuantStudio 7 Flex Real-Time PCR System. Relative mRNA expression was normalized to the housekeeping gene *L32*. The following primers were used:

> FOLR1-F: AGCACCACAAGGAAAAGCCAGG
>
> FOLR1-R: GTGCCATCTCTCCACAGTGGTT
>
> L32-F: ACAAAGCACATGCTGCCCAGTG
>
> L32-R: TTCCACGATGGCTTTGCGGTTC

### Protein isolation and Western blot

Cells were collected in ice-cold PBS and lysed in 1X RIPA buffer (Thermo Scientific, J63324.AK) and 1X Halt Protease and Phosphatase Inhibitor cocktail (Thermo Scientific, 78446). Lysates were centrifuged and supernatants were collected. Protein was resolved on 4-20% SDS-PAGE gels and transferred to nitrocellulose membranes. Membranes were blocked in 5% milk and incubated with primary antibodies followed by peroxidase-conjugated secondary antibodies. Membranes were developed using the Pierce™ ECL Western Blotting Substrate (ThermoFisher, 32209). The following antibodies were used: anti-Folate Binding Protein (EPR20277) (Abcam, ab221543, 1:5000), anti-Vinculin (Millipore Sigma, 05-386, 1:5000), anti-β-Actin (AC-15) (Sigma-Aldrich, A5441, 1:5000), anti-Phospho-p44/42 MAPK (T202/Y204) (CST, 9101S, 1:1000), anti-p44/42 MAPK (Erk1/2) (CST, 9102S, 1:1000), anti-Phospho-Histone H2A.X (Ser139) (CST, 2577, 1:1000), anti-Histone H2A.X (CST, 2595, 1:1000), anti-Histone H3 (D2B12) (CST, 4620S, 1:1000), anti-Mouse IgG (H+L), HRP-conjugate (Promega, W4021, 1:5000), and Goat anti-Rabbit IgG (H+L), HRP-conjugate (ThermoFisher, 31460, 1:5000).

### Generation of FOLR1 knockout cells

An sgRNA targeting FOLR1 (target sequence: 5’-AGGGTTTAACAAGTGCGCAGTGG-3’; forward oligo: CACCGAGGGTTTAACAAGTGCGCAG; reverse oligo: AAACCTGCGCACTTGTTAAACCCTC) and non-targeting control (forward oligo: CACCGGCGAGGTATTCGGCTCCGCG; reverse oligo: AAACCGCGGAGCCGAATACCTCGCC) were cloned into the lentiCRISPR v2 plasmid (Addgene #52961) using standard molecular cloning techniques. Lentiviral particles were produced by transfecting HEK293T cells with 1 mg/mL PEI. Viral supernatants were collected 48 hours post-transfection and filtered through a 0.45 µm filter. H1373 and HCC44 cells were seeded at 40-50% confluency and transduced with lentiviral supernatants supplemented with 4 µg/mL polybrene to enhance transduction efficiency. After 24 hours, the medium was replaced with fresh complete culture medium. Puromycin selection (6.4 µg/mL for H1373 and 2.6 µg/mL for HCC44) was initiated 48 hours post-transduction and continued for 5 days until all untransduced control cells were eliminated. Puromycin-resistant cells were screened for FOLR1 knockout by western blot and flow cytometry.

### Flow cytometry

Cells were harvested using TrypLE™ Express Enzyme (1X) (ThermoFisher, 12604021), washed with PBS, and resuspended in FACS buffer (PBS containing 1% FBS). For surface staining, cells were incubated with unconjugated or fluorophore-conjugated antibodies against the target protein for 30 minutes at 4°C protected from light. For unconjugated primary antibodies, cells were washed three times with FACS buffer and incubated with the appropriate fluorophore-conjugated secondary antibodies for 30 minutes at 4°C, protected from light. Following staining, cells were washed three times with FACS buffer and resuspended in staining buffer for acquisition. For viability assessment, cells were stained with SYTOX™ Blue Dead Cell Stain (ThermoFisher, S34857) according to the manufacturer’s instructions. Data was acquired on a Beckman Coulter CytoFLEX flow cytometer, with at least 10,000 events collected per sample. Data was analyzed using FlowJo software (version 10.10.0). Cells were gated sequentially on forward and side scatter to exclude debris, on singlets to exclude doublets, and on live cells prior to analysis of marker expression. Marker expression was quantified as mean fluorescence intensity (MFI) fold change and percentage of positive cells. The following antibodies were used: mirvetuximab (anti-FOLR1, MCE, HY-P99225, 10 µg/mL), anti-human IgG (MCE, HY-P990100, 10 µg/mL), gatipotuzumab (anti-TA-MUC1, MCE, HY-P99634, 10 µg/mL), PE anti-human TACSTD2 (TROP2) antibody (Biolegend, 363803, 2.5 µg/mL), Podocalyxin Monoclonal Antibody (B34D1.3), PE (eBioscience™/Invitrogen, 12-8873-42, 50 µg/mL), PE anti-human CD171 (L1CAM) (Biolegend, 371603, 2.5 µg/mL), Goat anti-Human IgG (H+L) Cross-Adsorbed Secondary Antibody, Alexa Fluor™ 647 (Invitrogen, A-21445, 1 µg/mL), PE anti-human IgG Fc (Biolegend, 410708, 2.5 µg/mL), and Placental Alkaline Phosphatase Monoclonal Antibody (H17E2) (Invitrogen, MA1-20245, 10 µg/mL).

### Internalization assays

Cells were seeded into 96-well plates the day before assay start (5,000 cells/well) to obtain 30%-40% confluency. Mirvetuximab (or control IgG) was conjugated with pHrodo antibody labeling kit (ThermoFisher, P36023) at a 1:3 molar ratio according to manufacturer’s protocol. The next day, the cells were treated with either DMSO or JDQ443/RMC7977 along with conjugated IgG or mirvetuximab antibodies. Imaging and analysis were performed using IncuCyte Zoom/S3 live-cell imagers (Essen Biosciences) and quantified using the IncuCyte software.

### In Vivo studies

All animal procedures were conducted under approved protocols aligned with guidelines from the Institutional Animal Care & Use Committee (IACUC) at Dana-Farber Cancer Institute (DFCI).

### Tumor implantation and tumor growth monitoring

Cell lines (H1373: 2.5 × 10^6^ cells, HCC44: 7.5 × 10^6^ cells) were implanted subcutaneously into the flank of 8-week-old female NU/J mice (Jackson Laboratory), and patient derived tumors were implanted in 8- to 10-week-old female NSG (NOD.Cg-Prkdcscid Il2rgtm1WjI/SzJ, Jackson Laboratory) mice, in PBS containing 50% BD matrigel. For efficacy studies, treatment was started when the average tumor size reached 200-300 mm^3^ and continued for 30 days. Tumor size was measured twice weekly by calipers and calculated using 0.5 × L × W^2^. JDQ443 was supplied by Novartis and formulated fresh weekly in 0.5% methylcellulose/0.1% Tween 80 in water and administered at 100 mg/kg/day p.o. Mirvetuximab soravtansine-gynx obtained from the DFCI Pharmacy was diluted in 5% dextrose and administered intravenously at 5 mg/kg once weekly for 2 weeks (H1373), 3 weeks (HCC44), or 4 weeks (PDX DFCI-456).

### Immunohistochemistry

For ECAD, Ki67 and yH2AX staining, formalin-fixed, paraffin-embedded (FFPE) tissue sections (4-5 µm thick) were baked at 60°C for 30 minutes. Slides were deparaffinized in sequential xylene baths, then transferred through sequential baths of 100%, 100%, 95%, and 75% ethanol for 5 minutes each, and rinsed in water. Heat-induced antigen retrieval was performed using citrate buffer (pH 6) (Agilent Technologies, S2369). Endogenous peroxidases were blocked using 3% hydrogen peroxide (Sigma-Aldrich, H1009-500) in TBS for 10 minutes. Slides were blocked with 5% normal goat serum in TBST for 1 hour at room temperature. Tissue sections were incubated with primary antibodies diluted in 5% goat serum overnight at 4°C in a humidified chamber. The next day, slides were washed with TBST and incubated with SignalStain® Boost IHC Detection Reagent (CST, 8114) for 30 minutes at room temperature. Signal detection was performed using DAB substrate (CST, 8059). Slides were washed with water to stop the reaction and then stained with Harris Hematoxylin (ThermoFisherScientific) for 1 minute and rinsed with warm tap water for 5 minutes. Slides were dipped eight times in 0.25% HCl in 70% ethanol and rinsed with tap water again for 5 minutes. Slides were dehydrated and mounted for analysis by bright-field microscopy. Slides were scanned using a PhenoImager™ HT (Akoya Biosciences) with the Phenochart 2.2.0 software.

For FOLR1 staining, immunohistochemistry was performed on the Leica Bond III automated staining platform using the Leica Biosystems Refine Detection Kit (Leica; DS9800). FFPE tissue sections were baked for 30 minutes at 60°C and deparaffinized (Leica AR9222) prior to staining. Primary antibodies were visualized via DAB, and counterstained with hematoxylin (Leica DS9800). The slides were rehydrated in graded alcohol and coverslipped using the HistoreCore Spectra CV mounting medium (Leica 3801733). Antibody FOLR1 from Abcam catalog number ab221543, clone EPR20277 was run at 1:1000 dilution with a 20M EDTA antigen retrieval (Leica ER2 AR9640), RAB DAB protocol used.

### Deep Mutational Scanning

G12C DMS library generation and lentivirus production was previously described in Awad et al.^8^ DMS for G12C resistance mutation was performed as described in Weller et al.^36^ with the following modification. Ba/F3 cells were spin-infected with the lentiviral supernatant of G12C DMS library at a multiplicity of infection (MOI) of 0.1 in the presence of diethylaminoethyl-dextran (DEAE-dextran, Sigma #D9885, 8 µg/mL). After 24 hours of transduction, cells were washed with fresh media then was selected in the presence of puromycin at 2 ug/ml for 72 hours. Live cells were washed divided and treated with either DMSO, JDQ443 (20nM) or sotorasib (800nM) for total 6 days with fresh media and drug replenishment on day 3.

### Whole exome sequencing sample collection

Genomic DNA was extracted using Qiagen AllPrep DNA/RNA Mini Kit (Qiagen; Cat. # 80204). WES was performed by Novogene and the library was prepared using Agilent V6 and the sequencing was carried out on a NovaSeq (Illumina, PE150).

### RNAseq sample collection

Cells were trypsinized and washed with PBS twice, cell pellets were collected, and total RNA was extracted using the RNeasy Plus Kits for RNA Isolation according to the manufacturer’s protocol (Qiagen; Cat. #73404). All the conditions and cell lines were prepared in triplicate. Tumors harvested from cell line derived xenografts were snap frozen in liquid nitrogen followed by tissue homogenization and RNA extraction using the RNeasy Plus Kits for RNA Isolation according to the manufacturer’s protocol (Qiagen; Cat. #73404). Total RNA was sent to Novogene for mRNA-seq library preparation and NGS using Illumina NovoSeq.

### Quantitative Proteomics

Cells were cultured in 10 cm dishes and treated for the indicated times. On the day of collection, cells were washed with ice-cold PBS after media was removed. Cell pellets were collected and lysed in 8 M urea in 200 mM EPPS pH 8.0 supplemented with Halt Protease and Phosphotease Inhibitor Cocktail. 50 μg of protein lysates from each condition were used for quantitative proteomics. Protein reduction and alkylation were performed with 10 mM TCEP and 20 mM IAA at 37°C for 45 minutes. Protein was clarified using the SP3 method: 10 μL SP3 beads and 60 μL methanol were added to each sample and rotated at 21°C, 10 RPM for 10 minutes. SP3 beads were washed twice with 80% v/v ethanol. 1 μg trypsin and 1 μg lysC in 50 μL EPPS (200 mM pH 8.0) were added into each sample and incubated at 37°C shaking at 1200 RPM for 16 hours. The next day, 5 μL TMTpro stock (20 mg/ml) and 15 μL ACN were added to each digest and incubated at 21°C 1200 RPM for 60 minutes. TMTpro reaction was quenched via 0.5% hydroxylamine at 21°C 1200 RPM for 10 minutes. 2 μL labeled samples were pooled in 200 μL 1% FA, then desalted via C18 Stage-Tip. Peptides were injected for a 45-minute gradient DDA hrMS2 acquisition and the data was used to determine peptide quantity in each TMTpro channel. Equal amount TMTpro-labeled peptides were pooled together and desalted via C18 Sep-Pak (60 mg). Peptides were eluted with 70%ACN+1%FA and dried via SpeedVac. Phophopeptides were isolated from dried peptides according to manufacturer’s protocol using Thermo Fe-NTA Phosphopeptide Enrichment Kit. Enriched phosph-peptides were desalted via C18 Stage-Tip and dried in SpeedVac. Phospho-peptides were injected into an Ortbitrap Eclipse Tribrid Mass Spectrometer (Thermo Fisher Scientific) coupled with a nano-LC at 120-minute gradient. Data was acquired in data-dependent acquisition (DDA) high-resolution MS2 (hrMS2) mode with FAIMS Pro Duo to minimize the TMTpro co-isolation issue. After phosphopeptide enrichment, the flow-through was dried in SpeedVac and then fractionated offline into 24 fractions through a basic pH reverse phase C18 column. These 24 fractions were desalted via C18 Stage Tips and the odd 12 fractions were analyzed in an Orbitrap Exploris 480 Mass Spectrometer (Thermo Fisher Scientific) coupled with a nano-LC and separated in a 90-minute gradient. MS acquisition was performed in DDA hrMS2 mode and peptide ions were pre-selected via FAIMS Pro Duo before MS inlet.

### Surfaceome enrichment

Surfaceome enrichment was performed as previously described with modifications^93^. Cells were cultured in two 15 cm dishes per condition (DMSO, 2 days, 7 days, parental and resistant) at least 2 days prior to labeling. At the time of collection, cells were washed with ice-cold PBS followed by incubation with 5 mL of ice-cold 1 mM EDTA in PBS on a shaker until cells began to detach, approximately 5-15 minutes depending on the cell line. Cells were collected by centrifugation, and cell pellet was suspended in 500 μL of surface protein labeling solution (1 mM sodium periodate, 200 µM aminooxy-biotin in PBS pH 6.5) and transferred to Eppendorf Protien LoBind 1.5 mL tubes. Next, 50 μL of aniline (1:10) were added and samples were incubated in the cold room for 30 minutes with rotation. After incubation, samples were centrifuged and transferred to 15 mL tubes and washed twice with ice-cold PBS. Cell pellets were then lysed with 500 μL 2X RIPA with Halt Protease Inhibitor at 4°C with rotation for 1 hour. After lysis samples were centrifuged at 20,000 g at 4°C for 15 minutes and the supernatant was collected. 100 μL streptavidin agarose beads (200 μL slurry) were washed 2X with PBS and added into each lysate, then incubated at 4°C 10 RPM for 16 hours. Next day, agarose beads were pelleted and washed 2X with 4 M Urea. Then beads were incubated with EPPS containing 10 mM TCEP for 30 minutes, followed by addition of 10 mM IAA and incubation for 30 minutes at room temperature. After reduction and alkylation, beads were washed 3X with MS-grade water. 1 μg trypsin and 1 μg LysC in 50 μL 200 mM EPPS (pH 8.0) were added into each sample, and the samples were incubated at 37°C 1200 RPM for 16 hours. Next day, 3 μL TMTpro stock (20 mg/ml) and 15 μL ACN were added into each sample, then incubated at 21°C 1200 RPM for 60 minutes. TMTpro labeled samples were quenched via 7 μL hydroxylamine stock (5%) at 21°C 1200 RPM for 10 minutes. Quenched samples were pooled together. 10% FA was added to acidify TMTpro-labeled samples and was desalted via C18 Sep-Pak (60 mg). Desalted peptides were eluted from Sep-Pak with 70% ACN and 1% FA and dried via SpeedVac. Dried peptides were dissolved in 10 μL 5%ACN and 5%FA and ready for MS acquisition. 10 μL peptides were analyzed in Exploris 480 coupling with a nano-LC separated in a 120-minute gradient. MS acquisition was performed in data-dependent (DDA) high-resolution MS2 mode and peptide ions were pre-selected via FAIMS Pro Duo before MS inlet.

### Data analysis

#### Mutation and copy number alteration analysis from bulk WES

Raw sequencing data were processed using standardized workflows^94^ implemented on the Terra platform. Paired-end FASTQ files were first converted to unmapped BAM (uBAM) format using the *paired-fastq-to-unmapped-bam* workflow. Subsequent preprocessing, including alignment to the hg38 reference genome, duplicate marking, and base quality score recalibration, was performed using the *Processing-for-variant-discovery-gatk4 pipeline*.

Putative somatic single-nucleotide variants (SNVs) and small insertions/deletions (indels) were identified using GATK Mutect2 through the mutect2-gatk4-human workflow (snapshot 21). Tumor-only variant calling was performed for all vehicle and resistant samples using a publicly available panel of normals (PoN; *gs://gatk-best-practices/somatic-hg38/1000g_pon.hg38.vcf.gz*) to suppress recurrent sequencing artifacts and technical noise. Filtered Mutect2 VCFs were converted to Mutation Annotation Format (MAF) using **vcf2maf v1.6.22** through a custom batch-processing script, with variant annotation performed using **Ensembl Variant Effect Predictor (VEP) v114.2** and the **release 114 GRCh38 cache**. Annotation was performed against the GRCh38/hg38 reference genome. For VAF analysis, PASS protein-altering small variants were retained, and VAFs were calculated primarily from tumor alternate and total read counts.

Copy-number alteration analysis was performed using a customized CNVkit-based workflow. CNVkit analysis used publicly available hg38 reference resources, including the access-region file *CNVkit_access-10kb.hg38.bed* and gene-annotation file *CNVKit_refFlat_hg38.txt*. For each evaluable cell-line pair, the resistant derivative was compared with its matched vehicle control to generate genome-wide relative copy-number profiles. Positive and negative log_2_ copy-number ratios represent relative gains and losses in resistant cells, respectively.

### Bulk RNA-seq Processing and Analysis

Reads were filtered and aligned using standard methods by Novogene, where reads were removed if they contained adapter contamination, unknown nucleotides more than 10 percent (N>10%), or low quality nucleotides (BQ < 5) constitute > 50% of the read. Reads were aligned with HISAT2^95^ and normalized for library size. Normalized RNA sequencing counts were log_2_-transformed for statistical evaluation. Differential gene expression analyses were conducted in R (version 4.5.3) utilizing the limma package^96^ (version 3.66.0) log2-normalized counts. To identify conserved transcriptional dynamics across the heterogeneous cell line panel, combined analyses were performed utilizing the duplicateCorrelation function^97^. This approach incorporated the individual cell line as a blocking factor within the linear mixed-effects model to account for basal transcriptional variance. Empirical Bayes moderation was subsequently applied across all contrasts to compute log_2_ fold-changes and adjusted *p*-values.

### Proteomics data processing

Raw mass spectra were analyzed using MaxQuant software^98^ (version 2.6.7.0) against the UniProt Human Reference Proteome database^99^ (accessed January 2025; 20,417 sequences), supplemented with common contaminants. Searches were configured for Trypsin/P digestion, permitting a maximum of three missed cleavages. Peptide lengths were constrained to 8–25 amino acids with a maximum mass threshold of 4,600 Da.

Identification required a minimum of one razor peptide, with the false discovery rate (FDR) strictly controlled at <0.01. For multiplexed quantification, MS2 reporter ions were extracted using a mass tolerance of 0.003 Da alongside manufacturer-provided TMTpro-18plex isotope correction factors. Reporter ion quantification was restricted to spectra possessing a precursor intensity fraction (PIF) strictly greater than 0.75. Cysteine carbamidomethylation was designated as a fixed modification. Variable modifications included methionine oxidation, protein N-terminal acetylation, and for the phosphoproteomic datasets, phosphorylation of serine, threonine, and tyrosine (STY). A maximum of five modifications per peptide was permitted, and default Orbitrap instrument parameters were applied.

### Normalization and differential expression

MaxQuant output tables were imported into R (version 4.5.3) for downstream processing and statistical evaluation. Phosphosite identifications were filtered to retain only high-confidence Class I sites demonstrating a localization probability > 0.75. Following the exclusion of reverse hits and known contaminants, reporter ion intensities were sum-normalized within their respective cell lines and TMT multiplex batches to account for technical variance. Differential expression for both global and surface-enriched proteomics was modeled using limma^96^ (version 3.66.0). To identify broadly conserved adaptive responses across the diverse NSCLC models, analytical models incorporated a cell line-specific blocking term to correct for inherent baseline expression differences^97^. Missing values were deliberately not imputed. Global variance was visualized via principal component analysis (PCA) using the factoextra^100^ R package, applying zero-filling for missing data and mean-centering. Sample similarities were quantified using Pearson’s correlation coefficients, and the resulting correlation matrices were hierarchically clustered applying the McQuitty linkage method.

### Database integration and metadata annotation

To evaluate gene essentiality, we queried the Cancer Dependency Map (DepMap, release 22Q2) using the AnnotationHub^101^ R package. The dataset was filtered to include only KRAS-mutant non-small cell lung cancer (NSCLC) cell lines based on the accompanying mutation calls (22Q2). For every gene, a median dependency score was computed across this KRAS-mutant NSCLC subset. Subcellular localizations were assigned using the COMPARTMENTS resource^102^. Each protein was categorized into primary cellular regions (Nucleus, Cytoplasm, Mitochondrion, Lysosome, Plasma membrane, Cytoskeleton, Endosome, Peroxisome, and Golgi apparatus). We utilized the highest confidence score for each gene to assign localizations, permitting individual proteins to map to multiple compartments. Additionally, regulatory phosphosites with established biological functions were identified by cross-referencing our dataset with the Regulatory Sites collection from PhosphoSitePlus^38^. To delineate the cellular surfaceome, we generated a unified index of plasma membrane-localized proteins by integrating annotations from two extensive repositories: the Cancer Surfaceome Atlas^42^ and the *in silico* human surfaceome database^41^. We manually annotated surface proteins corresponding to clinically investigated antibody-drug conjugates (ADCs) recently catalogued.

### Gene set enrichment analysis (GSEA) and pathway trajectories

Pathway enrichment was executed utilizing the fgsea^103^ and msigdbr^104^ packages in R. To capture both the magnitude and statistical significance of differential gene expression, transcripts were ordered for GSEA using a composite ranking metric: the log_2_(fold-change) multiplied by the min-max normalized -log_10_(adjusted *p*-value). Evaluated gene sets comprised canonical collections from the Molecular Signatures Database (MSigDB) alongside empirically derived, custom signatures for KRAS-ERK^105^ and MYC^105^ dependencies.

To model the longitudinal dynamics of these enriched networks, GSEA net enrichment scores (NES) across all experimental timepoints were grouped using Ward’s minimum variance hierarchical clustering. The optimal number of temporal clusters was empirically derived by the gap statistic technique implemented in the cluster^106^ R package. Because distinct biological pathways frequently share overlapping constituent genes, redundant pathway terms were systematically identified by calculating the Szymkiewicz-Simpson similarity index on the leading-edge gene subsets from the GSEA output. These overlapping pathways were subsequently consolidated using Ward’s hierarchical clustering to define distinct, non-redundant trajectory modules.

### Kinase motif profiling and network analyses

Differentially regulated phosphosites were assessed for overrepresented kinase motifs using the human serine/threonine kinome atlas^39^. Relative kinase activities were estimated by calculating the product of the log_2_ enrichment factor and the min-max normalized adjusted *p*-value corresponding to the most significant direction of enrichment for each respective kinase. Kinase-substrate interaction networks were generated as described previously^40^. Briefly, an aggregate interaction score was calculated for each phosphoprotein by combining three weighted lines of evidence: (1) a kinase-substrate score derived by preferential phosphosite motif specificities, which synthesizes kinase preferences for individual phosphosites using the dataset-specific kinase activity measurements; (2) curated, empirically validated kinase-phosphosite interactions extracted from the PhosphoSitePlus^107^ database; and (3) documented kinase-substrate associations obtained from the PhosphoNetworks and BioGRID (v4.4.213)^108^ repositories. To isolate and visualize only the most highly confident signaling connections, the final plotted networks were restricted to interactions possessing an absolute composite score exceeding one standard deviation.

### Integration of RNAseq, global proteomics, and surface-enriched proteomics

Differential expression profiles across our RNA-sequencing, global proteomics, and surface-enriched proteomics datasets were integrated by utilizing Stouffer’s weighted Z-score method. For each target, modality-specific z-scores were computed and assigned a mathematical sign to retain the direction of regulation (i.e., up- or downregulation). These signed z-scores were subsequently weighted by the square root of their respective sample sizes and aggregated. The summed values were then normalized against the square root of the sum of the squared weights, producing a composite Z-score that conforms to a standard normal distribution under the null hypothesis. Finally, these integrated Z-scores were translated into two-sided p-values via the standard normal cumulative distribution function.

### Feature engineering for machine learning

Physiological baseline features were derived from the Human Protein Atlas (HPA) consensus tissue dataset^43^, where normal tissue transcript abundance was calculated as log_2_(TPM + 1). Placental tissue was excluded from these baseline calculations. Normal tissue immunohistochemistry (IHC) profiles from the HPA were converted into a normal IHC score, quantified as the fraction of profiled healthy tissues exhibiting moderate to strong protein detection. Clinical lung adenocarcinoma profiles were established using tumor-specific protein prevalence derived from HPA lung adenocarcinoma IHC data, scored as the proportion of evaluated patient tumors demonstrating detectable staining. Prognostic impact in lung adenocarcinoma was determined from HPA survival analyses^44^ and binarized, assigning a value of 1 for significant associations with patient survival at a log-rank *P* < 0.0001, and 0 otherwise. Patient tumor transcript levels were quantified as the median log_2_(TPM + 1) across the TCGA-LUAD cohort. Additionally, tumor-versus-normal protein-level alterations were acquired from the CPTAC lung adenocarcinoma dataset and parameterized as the relative protein log_2_ fold-change in primary tumors versus adjacent normal tissue. Treatment-induced experimental adaptations were derived directly from our matched JDQ443-treated KRAS^G12C^-mutant NSCLC models across RNA-sequencing, whole-cell global proteomics, and surface-enriched proteomics. For each platform, differential expression was modeled across the pooled cell line cohort. To isolate consistent on-target adaptations while mitigating assay dimensionality, expression changes were summarized as the median log2 fold-change across the acute and sustained timepoints (2 days and 7 days), excluding the acquired resistance state. This yielded three coordinated experimental features: median transcriptomic change, median global proteomic change, and median surface-enriched proteomic change. Finally, transcripts were annotated for plasma membrane localization based on consensus surfaceome membership and cataloged for current clinical ADC status.

### Ridge logistic regression training and target scoring

The training set was defined by the following criteria and defined as the training set with Class = 1: the target needed to have a clinically investigated ADC as defined above, the target required a median log_2_ fold-change > 0.5 across the 2-day and 7-day treatments in at least one experimental platform (RNA-seq, global proteomics, or surfaceome proteomics), the target had to demonstrate robust baseline overexpression in primary lung tumors relative to healthy organs defined by an expression ratio greater than 1 when comparing median tumor transcript abundance in TCGA-LUAD to median expression across normal HPA tissues. All other evaluated genes were categorized as the background comparison class (Class = 0). This non-positive label denotes the absence of a qualifying, clinically investigated ADC fulfilling these criteria rather than confirmed biological intractability. A ridge-penalized logistic regression model (*α* = 0) was utilized by the glmnet R package to compute a continuous ADC target-prioritization score. Ridge regularization was explicitly selected to shrink correlated features, such as tumor RNA and tumor IHC, toward zero without eliminating pre-specified biological predictors, thereby minimizing overfitting. The 10-feature matrix for genes with complete cases was standardized to zero mean and unit variance, and the optimal regularization parameter (*λ*) was selected by five-fold cross-validation using binomial deviance, utilizing the conservative *λ*_1*se*_ value. To score the full candidate gene set, out-of-sample feature vectors were standardized using the centering and scaling parameters derived from the complete-case training matrix, with any missing values zero-filled. The model-generated linear predictor (*LP*), representing the log-odds score, was calculated for each gene. To provide an intuitive, relative prioritization metric, scores were min-max normalized and transformed to a (-1,1) scale.

### TCGA transcriptome and CPTAC proteome analysis

To evaluate the concordance between FOLR1 transcript abundance and protein expression in primary human tumors, matched proteomic and transcriptomic profiles from lung adenocarcinoma patients were analyzed from the CPTAC BCM-LUAD cohort. Analysis was restricted strictly to primary tumor specimens, excluding adjacent normal tissues. Normalized FOLR1 protein abundance (log_2_ reference intensity normalized) and corresponding RNA expression (RSEM upper-quartile log_2_(x + 1)) were integrated across matched patient identifiers.

### Clinical cohort processing, stratification, and differential expression

Transcriptomic profiles from patients with KRAS-mutant lung adenocarcinoma were obtained from The Cancer Genome Atlas (TCGA-LUAD), the CodeBreaK 100/200 trials (GSE295032), and the KRYSTAL-1 trial (PRJNA591860). For TCGA-LUAD, raw count matrices were filtered to retain protein-coding transcripts with greater than 20 reads across at least 75% of samples, followed by trimmed mean of M-values (TMM) normalization via edgeR to compute counts per million (CPM). Patients harboring KRAS^G12C^ mutations were stratified into FOLR1-high and FOLR1-low subsets based on a median split of FOLR1 CPM. Count distributions were transformed using voom, and differential gene expression between FOLR1-high and FOLR1-low tumors was evaluated using linear modeling and empirical Bayes moderation (eBayes) in limma (version 3.66.0). For the CodeBreaK 100/200 cohort, individual sample files provided as transcripts per million (TPM) were concatenated into a unified expression matrix. Patients were bifurcated into FOLR1-high and FOLR1-low groups by the median FOLR1 TPM, and TPM values were log_2_-transformed with a pseudocount offset (log_2_(TPM + 1)). Differential expression between groups was modeled directly using limma with mean-variance trend adjustment (trend = TRUE). For the KRYSTAL-1 cohort, raw read counts were converted to TPM by first dividing read counts by transcript length in kilobases to determine reads per kilobase (RPK), followed by scaling by the sum of sample RPK. Patients were divided into FOLR1-high and FOLR1-low subsets using a median split of FOLR1 TPM, and differential expression on raw counts was modeled using voom transformation and empirical Bayes moderated linear models in limma

### Gene set enrichment analyses of patient cohorts (GSEA and ssGSEA)

To assess biological pathways associated with FOLR1 expression status within each clinical cohort, genes were ranked by their log_2_ fold-change derived from the FOLR1-high versus FOLR1-low differential expression comparisons. Gene set enrichment analysis was executed using fgseaSimple (10,000 permutations) against canonical MSigDB collections (Hallmark, Reactome, and KEGG) supplemented with empirically validated KRAS-ERK^105^ and MYC^105^ response signatures. To quantify pathway activity at single-patient resolution, single-sample GSEA (ssGSEA) was performed across combined patient profiles from the CodeBreaK and KRYSTAL-1 trials. Log_2_-transformed TPM expression matrices (log_2_(TPM + 1)) were mapped to common gene symbols and evaluated using the ssgseaParam implementation within the GSVA R package, normalizing enrichment scores to a [−1, +1] scale. Evaluated signatures included established alveolar type 1 (AT1), alveolar type 2 (AT2)^54^, and KRAS transcriptional programs, alongside in-house KRAS-dependent gene sets. Pairwise associations between single-patient ssGSEA signature scores and log_2_-transformed FOLR1 transcript levels were quantified using Pearson correlation coefficients and linear regression models.

### DMS analysis methods

DMS data were processed as previously described^8,36^. Briefly, reads were aligned to the KRAS^G12C^ reference sequence and collapsed to reads per codon at each position for each amino acid substitution included in the library, using AnalyzeSaturationMutagenesis v1.0 (ASMv1.0), software developed at the Broad Institute^109^. Substitutions at position S65, E98, and T158, were removed from the dataset as these positions were not covered by the library and therefore, variants at these positions likely represented sequencing and/or PCR artifacts. Additionally, redundant codons encoding the same amino acid substitution were averaged. The abundance of each allelic variant was calculated as the fraction of its reads relative to the total library-wide reads in each sample independently. Variant abundances were defined as the log_2_ fold change (Log_2_FC) between JDQ443-treated and DMSO-treated samples and were computed by applying a moderated t-test with empirical Bayes variance shrinkage as implemented in the R package limma (version 3.54.2).

Alleles were called enriched if Log_2_FC ≥ 1 and Benjamini-Hochberg (BH) adjusted p-value < 0.05. These per-allele enrichment calls were then used to compute per-residue enrichment scores via Fisher’s exact test against the background of all other codons, with a Haldane pseudocount applied to the odds ratio calculation. Enrichment values, including the average Log_2_FC and number of enriched alleles per residue, were then mapped onto the crystal structure of KRAS^G12C^-GDP in complex with JDQ443 (PDB: 7R0M)^110^ using ChimeraX (version 1.11.1).

## Data availability

Mass spectrometry proteomics data have been deposited to the ProteomeXchange Consortium via the PRIDE partner repository with the dataset identifier PXD069016. Whole exome data is available on request. The RNA-seq data discussed in this manuscript have been deposited in NCBI’s Gene Expression Omnibus and are accessible through GEO Series accession number GSE347691

## Acknowledgements

We thank Andreas Weiss and Hans Voshol from Novartisvfor providing JDQ443 for in vitro and in vivo studies and expert advice. We thank the Dana-Farber/Harvard Cancer Center in Boston, MA, for the use of the Specialized Histopathology Core, which provided histology and immunohistochemistry service. Dana-Farber/Harvard Cancer Center is supported in part by an NCI Cancer Center Support Grant # NIH 5 P30 CA06516. We thank Prafulla Gokhale and Bamee Chayawatto of the Robert and Renee Belfer Center for Applied Cancer Science (Dana-Farber Cancer Institute) for generating and maintaining the patient-derived xenograft model and assistance with in vivo dosing and tumor monitoring. J.A. Paulo is funded by R35 GM156406. C.A.S was supported by NCI P50CA196510. This project was supported by a Dana-Farber/Novartis Drug Discovery and Translational Research Program (DDTRP) grant to A.J. Aguirre and J.D. Mancias. A.J. Aguirre is also funded by the Pancreatic Cancer Action Network and NIH R01 CA276268. J.D. Mancias is also funded by NIH R01 CA301542.

## Authors’ Contributions

A. Padhye Conceptualization, formal analysis, data curation, investigation, visualization, writing–original draft, writing–review and editing, supervision, methodology, project administration.

K. Kang Investigation, validation, writing–review and editing.

F. Kong Investigation, validation, writing–review and editing.

N. Sindoni Investigation, validation, writing–review and editing.

Q. Yu Conceptualization, investigation, visualization, supervision, methodology.

B. Jiang Formal analysis, data curation, investigation, visualization, writing–review and editing.

W. Chang Formal analysis, investigation, writing–review and editing.

O. Coker Formal analysis, investigation, writing–review and editing.

E. Kim Conceptualization, formal analysis, data curation, investigation.

J. Dilly Formal analysis, data curation, visualization, investigation, writing–review and editing.

M. Shin Investigation, methodology, writing–review and editing.

J.A. Paulo Investigation, methodology, writing–review and editing.

K. Sprouffske Formal analysis, data curation, investigation, writing–review and editing.

S.M. Brachmann Conceptualization, investigation, writing–review and editing, supervision, project administration.

C.A. Stalnecker Conceptualization, formal analysis, data curation, investigation, visualization, writing–original draft, writing–review and editing, supervision, methodology, project administration, funding acquisition.

A.J. Aguirre Conceptualization, formal analysis, data curation, investigation, visualization, writing–original draft, writing–review and editing, supervision, methodology, project administration, funding acquisition.

J.D. Mancias Conceptualization, formal analysis, data curation, investigation, visualization, writing–original draft, writing–review and editing, supervision, methodology, project administration, funding acquisition.

## SUPPLEMENTARY FIGURE LEGENDS

**Supplementary Figure S1. Characterization of KRAS^G12C^ models treated with KRASi.**

**A.** Western blot analysis of lung cancer cell lines treated JDQ443 (100 nM) for indicated timepoints. **B.** Dose response curves for isogenic pairs of JDQ443 resistant cell lines treated with JDQ443 for 5 days. Cell viability was determined by CellTiter-Glo assay and normalized to DMSO. Data represented as mean ± SD. **C.** Area under the curve analysis of curves in (**B)** using Graphpad PRISM. Statistical significance determined by one-way ANOVA. **D.** Western blot analysis of isogenic pairs of JDQ443 resistant cell lines for indicated timepoints. **E.** Tumor growth curves for H1373 xenografts treated with vehicle or JDQ443 (100 mg/kg) for 5, 14 or 45 days. Data are represented as mean tumor volume ± SEM. Statistical significance was determined by two-way ANOVA with Sidak’s multiple comparisons test. **F.** Variant allele frequency (VAF) trajectories of selected small variants in vehicle and resistant cell-line pairs. All retained protein-altering KRAS variants were displayed; for other prespecified cancer-relevant genes, one representative protein-altering variant per gene and cell line was selected, prioritizing the variant with the highest VAF in the resistant derivative. VAF is shown as a percentage on a logarithmic scale. Only PASS variants were plotted. Variants with a retained PASS call in only one condition are shown as a single point without imputation of the missing value or a connecting line; detected VAFs below 0.05% are displayed at the 0.05% plotting floor. An asterisk (*) in a protein-change label denotes a termination (stop) codon. **G.** Genome-wide CNVkit copy-number profiles of resistant derivatives relative to their matched vehicle controls. Grey points indicate bin-level CNVkit log_2_ copy-number ratios relative to the matched vehicle, and black lines indicate segmented copy-number profiles. Positive and negative values indicate relative copy-number gains and losses in resistant cells, respectively. Red dotted lines and labels mark the genomic positions of selected cancer-relevant genes. Significance for all panels: *p < 0.05, **p < 0.01, ***p < 0.001, ****p < 0.0001.

**Supplementary Figure S2. Characterization of the on-target mutational landscape of resistance to JDQ443 via deep mutational scanning.**

**A.** Schematic of the deep mutational scanning screen in a KRAS^G12C^ backbone conducted in the Ba/F3 cell line treated with the indicated regimens. Triplicates were harvested and sequenced for each timepoints. **B.** Heatmap depicting allelic enrichment of JDQ443 treatment relative to DMSO at each position in the KRAS^G12C^ protein, from N- to C-terminus. **C.** Left: Volcano plot depicting the allelic enrichment of JDQ443 treatment relative to DMSO. Resistance-associated alleles are indicated by the rectangle and were further used to define residues systematically driving resistance to JDQ443. Right: Residue-level enrichment for JDQ443 resistance-associated alleles. OR: Odds ratio. **D.** Structural mapping of residue-level resistance potential onto the KRAS^G12C-^GDP-JDQ443 crystal structure (PDB: 7R0M). Color indicates the average residue-level enrichment score, and the residue size is proportional to the number of substitutions at that position conferring resistance to JDQ443. **E.** Heatmap depicting allelic enrichment of sotorasib treatment relative to DMSO at each position in the KRAS^G12C^ protein, from N- to C-terminus. **F.** Residue-level enrichment for JDQ443 resistance-associated alleles (x -axis) and sotorasib (y-axis).

**Supplementary Figure S3. Global proteome, transcriptome and phosphoproteome analyses of KRAS^G12C^ mutant lung cancer models treated with KRASi.**

**A.** Differential expression and significance of proteomics data from H1373 xenografts following 5 days, 14 days, or 45 days of JDQ443 treatment. Volcano plots display the - log_10_ (p value adjusted) versus the median log_2_ fold-change of the relative protein abundance of KRASi-treated to vehicle-treated samples. The number of statistically significant upregulated (red, adjusted p value < 0.05, log_2_(fold-change) > 0.25), downregulated (blue, adjusted p value < 0.05, log_2_(fold-change) < -0.25), or not significant (grey, adjusted p value > 0.05) proteins are shown. Essential proteins (Chronos < -0.75) and proteins within KRAS-ERK regulated gene sets are shown in the bar plot (below). Significantly upregulated (red, adjusted p value < 0.05, log_2_(fold-change) > 0.25), downregulated (blue, adjusted p value < 0.05, log_2_(fold-change) < -0.25), or not significant (grey, adjusted p value > 0.05) are annotated. **B-C.** Unsupervised clustering of Pearson’s correlation analyses (top) and PCA (bottom) of expression changes across in vitro and in vivo proteome (**B**) and transcriptome (**C**) samples. **D.** Number of differentially expressed (|log_2_(fold-change)| > 0.25, adj.p val. < 0.05) essential proteins (top) or genes (bottom, |log_2_(fold-change)| > 0.5, adj. p val. < 0.05). **E.** Upset plot representing overlapping and non-overlapping detected proteins by proteomics across samples, showing the top 30 combinations and complete detected set size (left). **E.** Expression changes of top 100 up- and downregulated proteins for each time point from proteomics data across cell lines. Proteins not detected in a sample are represented by grey boxes.

**Supplementary Figure S4. Pathway enrichment across KRAS^G12C^ mutant in vitro and in vivo lung cancer models treated with KRASi.**

**A.** GSEA analysis of all in vitro and in vivo models for KEGG and Reactome gene sets. **B.** Heatmap showing the similarity index of top 50 significantly enriched or depleted gene sets from GSEA of transcriptomic data (top), as shown in Figure 1D, determined by Szymkiewicz-Simpson method using the leading-edge genes (bottom). Four distinct clusters identified by K-means clustering are highlighted and color corresponds to the gene set cluster as shown in Figure 1D.

**Supplementary Figure S5. KRASi treatment alters the phosphoproteome and kinase activity landscape in KRAS^G12C^ mutant lung cancer models.**

**A.** Volcano plots showing differential regulation of phosphosites for all cell lines analyzed together as biological replicates. All significantly downregulated (blue, log_2_(fold-change) < -0.5, adj.p val. <0.05) and upregulated (red, log_2_(fold-change) > 0.5, adj.p val.<0.05); known regulatory phosphosites are annotated in the barcode plot (bottom), and the top differentially expressed regulatory sites are labeled. Total number of significantly downregulated (blue text), upregulated (red text), or not significantly dysregulated phosphosites (grey text) is shown. **B.** Boxplot showing the number of significantly differentially expressed regulatory phosphosites (|log_2_(fold-change)| > 0.5, adj. p val. < 0.05) for all cell lines and timepoints. **C.** Volcano plot of differentially regulated kinases determined by motif profiling of phosphoproteomics data using differential expression of all cell lines pooled together as biological replicates.

**Supplementary Figure S6. Integrated analyses of surfaceome, proteome and transcriptome identify ADC targets upregulated with KRASi treatment in lung cancer.**

**A.** Correlation plots of surface-annotated proteins from surface-enriched proteomics versus global proteomics (left) and global proteomics versus transcriptomics (right) from all cell models analyzed together using cell lines as biological replicates for each time point. Genes detected in both datasets are shown in green, and genes detected in one dataset but not the other are shown on their corresponding x- or y-axes on the top or right of the correlation plot (margin). Margin plots are labeled either surfaceome proteomics only, global proteomics only, or transcriptome only. The number of significantly (adjusted p value < 0.05) upregulated (red, log_2_(fold-change) > 0.5), downregulated (blue, log_2_(fold-change) < -0.5), or non-significantly (adjusted p value > 0.05) expressed proteins detected in only one dataset shown as an inset on margin plots. **B**. Venn diagrams showing the number of surface-annotated proteins in surface-enriched proteomic, global proteomic, or transcriptomic datasets in HCC44 cells. **C**. Volcano plots of surface-annotated proteins in HCC44 cells treated for 2 days, 7 days, or at resistance to JDQ443. Known clinical ADC targets are annotated (bottom), and significantly (adjusted p value < 0.05) upregulated (green, log_2_(fold-change) > 0.5), downregulated (purple, log_2_(fold-change) < -0.5), or not significantly (grey, adjusted p value > 0.05) differentially regulated are shown or labeled on the volcano plot. **D**. A line plot displaying the coefficient weights (x-axis) assigned to each multi-omic feature (y-axis) utilized to train the machine learning model. Features driving a higher “ADC score” exhibit positive coefficients; these include surface annotation, experimental KRASi-induced upregulation (surfaceome proteomics log_2_FC, global proteomics log_2_FC, and RNAseq log_2_FC), and tumor-enriched expression metrics (lung cancer IHC score, TCGA LUAD RNA expression, and CPTAC LUAD tumor-normal log_2_FC). Conversely, features representing HPA LUAD prognostic factor and normal tissue expression (normal tissue IHC score and normal tissue RNA expression) exhibit negative coefficients.

**Supplementary Figure S7. FOLR1 expression is associated with distinct transcriptional programs across KRAS mutant lung cancer patients.**

**A.** Scatter plot detailing the correlation between FOLR1 RNA expression (RSEM UQ log_2_ (x+1)) and FOLR1 protein expression (log_2_ reference intensity normalized) across matched patient tumor samples. Transcriptomic and proteomic data were acquired from the BCM LUAD dataset via the Clinical Proteomic Tumor Analysis Consortium (CPTAC), with the analysis explicitly excluding normal adjacent tissues to focus solely on tumor-specific expression. **B-C.** RNA expression (log_2_(TPM + 1)) of FOLR1 in patient samples from KRYSTAL-1 trial (**B**) and CodeBreaK100/200 trial (**C**). **D.** GSEA of differentially regulated genes between FOLR1-high and FOLR1-low expressing samples from TCGA, KRYSTAL-1 trial, or CodeBreaK trial using REACTOME and KEGG datasets. **E.** Top three genes positively or negatively correlated with FOLR1 expression in patient samples from KRYSTAL-1 and CodeBreaK trial.

**Supplementary Figure S8. FOLR1 is upregulated in response to KRASi.**

**A.** FOLR1 log_2_(TPM+1) expression data from DepMap in KRAS mutant cancer cell lines. **B.** Percent positive cells upon JDQ443 (100 nM) at 2 days determined by flow cytometry. Data represented as mean ± SD. Statistical significance was determined by two-way ANOVA with Bonferroni’s multiple comparisons test. **C.** Volcano plots for RNAseq on H1373 and HCC44 cell lines treated with JDQ443 (100 nM) for 2 days. **D.** Drug wash-off experiment. HCC44 cells treated with JDQ443 (100 nM) for (i) 2, 4, or 9 days continuously, (ii) 7 days followed by 2 days without drug, or (iii) 7 days followed by 7 days without drug. FOLR1 expression determined by flow cytometry. Data represented as mean ± SD. Statistical significance was determined by ordinary one-way ANOVA. **E.** Wash-off experiment as in (**D**), with FOLR1 expression determined by RT-PCR. Data represented as mean ± SD. Statistical significance was determined by ordinary one-way ANOVA. **F.** Dose response curves for KRAS mutant ovarian cancer cell lines treated with a range of RMC7977 concentrations for 5 days. Cell viability was determined by CellTiter-Glo assay and normalized to DMSO. Data represented as mean ± SD. **G.** MFI fold change in FOLR1 upon RMC7977 (10 nM) treatment in ovarian cancer cells for 2 days. Data represented as mean ± SD. Statistical significance was determined by two-way ANOVA with Bonferroni’s multiple comparisons test. **H.** RT-PCR for FOLR1 expression in ovarian cancer cell lines at 2 days. Data represented as mean ± SD. Statistical significance was determined by two-way ANOVA with Bonferroni’s multiple comparisons test. **I-J.** Analysis of publicly available datasets for KRAS and EGFR mutant NSCLC treated with KRASi or EGFRi. Janes et al^111^, Solanki et al^112^, Li et al^10^, Salmon et al^113^, Nokin et al^114^, Criscione et al^115^, Gurule et al^116^, Moghal et al^117^. Significance for all panels: *p < 0.05, **p < 0.01, ***p < 0.001, ****p < 0.0001.

**Supplementary Figure S9. KRASi-induced FOLR1 upregulation does not mediate resistance to KRASi**

**A.** Representative histograms for flow cytometry analysis for FOLR1 in H1373 and HCC44 control and FOLR1 knockout cells. **B.** Dose response curves for control and FOLR1 knockout cells treated with JDQ443 for 5 days. Cell viability was determined by CellTiter-Glo assay and normalized to DMSO. Data represented as mean ± SD. **C.** MFI fold change in FOLR1 for cells cultured in different concentrations of folate media as indicated and treated with JDQ443 (100 nM) for 2 days. Data represented as mean ± SD. Statistical significance was determined by two-way ANOVA with Bonferroni’s multiple comparison tests. Significance for all panels: *p < 0.05, **p < 0.01, ***p < 0.001, ****p < 0.0001.

**Supplementary Figure S10. Combination of KRASi and mirvetuximab soravtansine-gynx shows efficacy in vitro.**

**A.** Dose response curve for KRAS mutant lung and ovarian cancer cell lines to DM4 for 2 days. Cell viability was determined by CellTiter-Glo assay and normalized to DMSO. Data represented as mean ± SD. **B.** Spearman correlation of antibody binding capacity (ABC) and area under the curve (AUC) for DM4 (calculated in GraphPad PRISM). **C.** Internalization assay of indicated antibodies conjugated with pHrodo, assessed at indicated timepoints for H1373, HCC44 and OVCAR5 cells treated with DMSO or KRASi determined by Incucyte. Data represented as mean ± SD. Statistical significance was determined using two-way ANOVA. **D.** Synergy assay for RMC7977 and mirvetuximab soravtansine-gynx (MIRV) across a range of doses for H1373 and HCC44 cells. Synergy score calculated by the HSA (highest single agent) method. **E.** Western blot for H1373 and HCC44 cells treated with DMSO, JDQ443 (100 nM), MIRV (5 nM), or combination for 24 hours and probed for indicated markers. **F.** Incucyte-based growth assay for H23 and H2030 cells treated with DMSO, JDQ443 (100 nM), MIRV (5 nM), or the combination over 5 days. Data represented as mean ± SD. Statistical significance was determined by two-way ANOVA. **G.** Synergy assay for JDQ443 and MIRV across a range of doses in H23 and H2030 cells. Synergy score calculated by the HSA (highest single agent) method. Significance for all panels: *p < 0.05, **p < 0.01, ***p < 0.001, ****p < 0.0001.

**Supplementary Figure S11. Combination of KRASi and mirvetuximab soravtansine-gynx shows efficacy in vivo.**

**A.** Individual tumor volume trajectories over time for mice bearing H1373 xenografts treated with vehicle (n = 8), mirvetuximab soravtansine-gynx (MIRV) (5 mg/kg, n = 8), JDQ443 (100 mg/kg, n = 9), concomitant combination (JDQ443 and MIRV initiated on day 0, n = 8), or sequential combination (MIRV initiated 5 days after JDQ443 treatment start, n = 10), shown as separate panels per arm (n = 8-10 per arm). Each line represents an individual animal. Grey boxes indicate the dosing period, and the dashed horizontal line marks the average tumor volume (200 mm³) at treatment start. **B.** Kaplan Meier survival curves for mice bearing H1373 xenografts treated with vehicle (n = 8), MIRV (5 mg/kg, n = 8), JDQ443 (100 mg/kg, n = 9), concomitant combination (JDQ443 and MIRV initiated on day 0, n = 8), or sequential combination (MIRV initiated 5 days after JDQ443 treatment start, n = 10). Median time to survival was 22.5, 26, 64, 91.5 and 94.5 days for vehicle, MIRV, JDQ443, concomitant, and sequential arms, respectively. Statistical significance was determined by log-rank test (overall p < 0.0001) with Holm-adjusted pairwise comparisons (***p < 0.001, vehicle vs. MIRV , vehicle vs. JDQ443, vehicle vs. concomitant; ****p < 0.0001, vehicle vs. sequential). **C.** Individual tumor volume trajectories over time for mice bearing HCC44 xenografts treated with vehicle (n = 5), MIRV (5 mg/kg, n = 7), JDQ443 (100 mg/kg, n = 5), or combination (n = 7), shown as separate panels per arm. Each line represents an individual animal. Grey boxes indicate the dosing period, and the dashed horizontal line marks the average tumor volume (200 mm³) at treatment start. **D.** Kaplan Meier survival curves for mice bearing HCC44 xenografts treated with treated with vehicle (n = 5), MIRV (5 mg/kg, n = 7), JDQ443 (100 mg/kg, n = 5), or combination (n = 7). Median time to survival was 45, 50, 45, and 120 days for vehicle, MIRV, JDQ443, and combination arms, respectively. Statistical significance was determined by log-rank test (overall p < 0.0001) with Holm-adjusted pairwise comparisons (**p < 0.01, vehicle vs. combination, JDQ443 vs. combination, MIRV vs. combination). **E.** Tumor growth curves for mice bearing DFCI-456 PDX (n = 5 per arm) treated with vehicle, MIRV (5 mg/kg), JDQ443 (100 mg/kg) or combination. Data are represented as mean tumor volume ± SEM. **F.** Kaplan Meier survival curves for mice bearing DFCI-456 PDX treated with treated with vehicle (n = 5), MIRV (5 mg/kg, n = 5), JDQ443 (100 mg/kg, n = 5), or combination (n = 5). Median time to survival was 57, 155, 64, and 159 days for vehicle, MIRV, JDQ443, and combination arms, respectively. Statistical significance was determined by log-rank test (overall p < 0.0001) with Holm-adjusted pairwise comparisons (**p < 0.01, vehicle vs. MIRV, vehicle vs. combination, JDQ443 vs. combination). **G.** Body weight (g) over the course of treatment for mice bearing H1373, HCC44, or DFCI-456 PDX xenografts, shown as mean ± SEM per treatment arm within each model. Significance for all panels: *p < 0.05, **p < 0.01, ***p < 0.001, ****p < 0.0001.

### Supplementary Tables

Supplemental Table S1: Metadata for proteomics searches

Supplemental Table S2: Differential expression of global proteomics

Supplemental Table S3: Differential expression of RNAseq

Supplemental Table S4: Gene set enrichment results for global proteomics data

Supplemental Table S5: Gene set enrichment results for RNAseq data

Supplemental Table S6: Differential expression results for phosphoproteomics data

Supplemental Table S7: Kinase enrichment analyses

Supplemental Table S8: Differential expression results for surface-enriched proteomics

Supplemental Table S8: Integrated proteomics and surface-enriched proteomics datasets

Supplemental Table S10: Features used to train ridge logistic model

