## Supplementary figures and images for "KRAS^G12C^ Inhibition Remodels the Non-Small Cell Lung Cancer Surfaceome to Uncover Targetable Vulnerabilities"

### Supplementary Figures

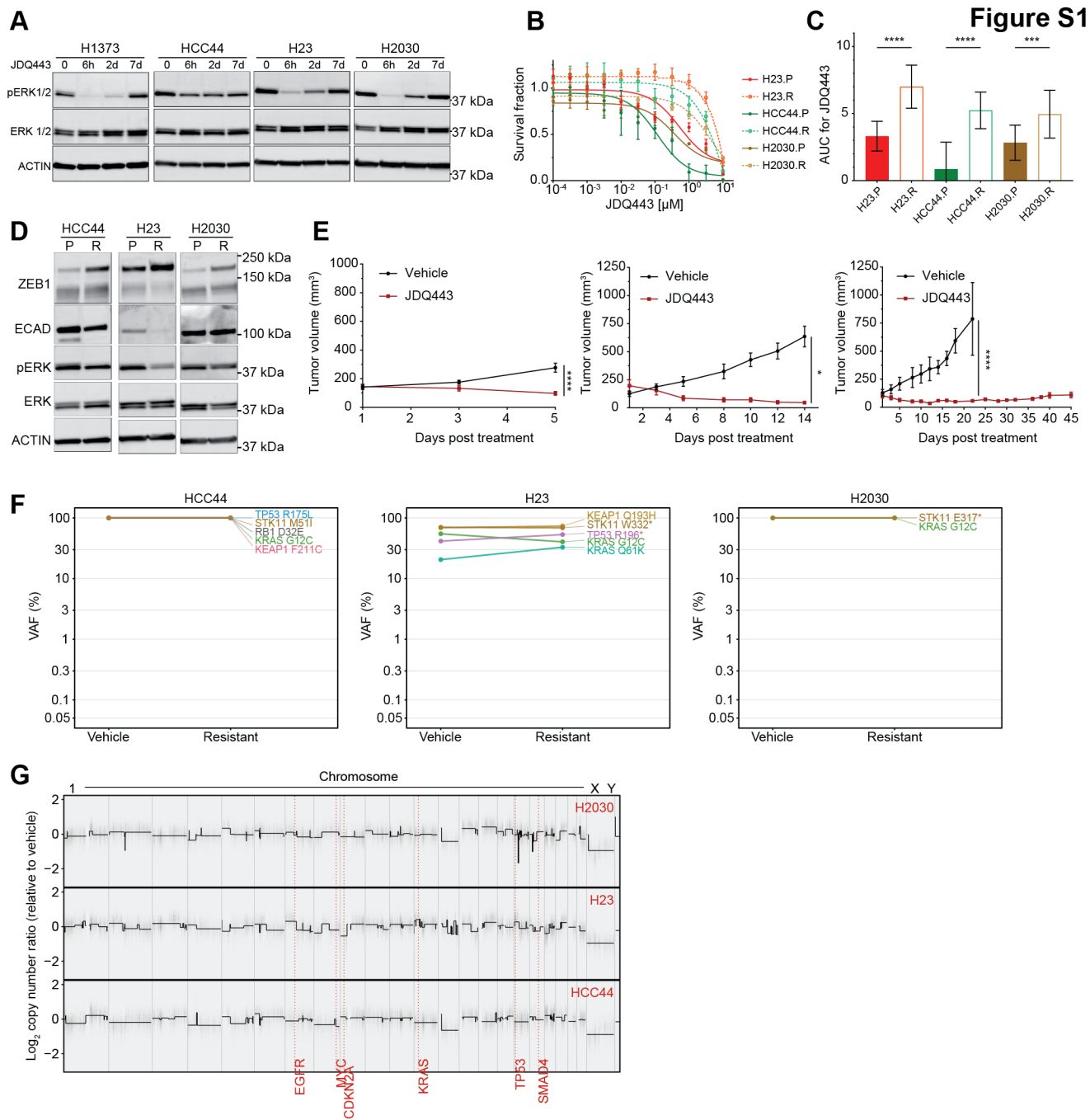

**Figure S2**

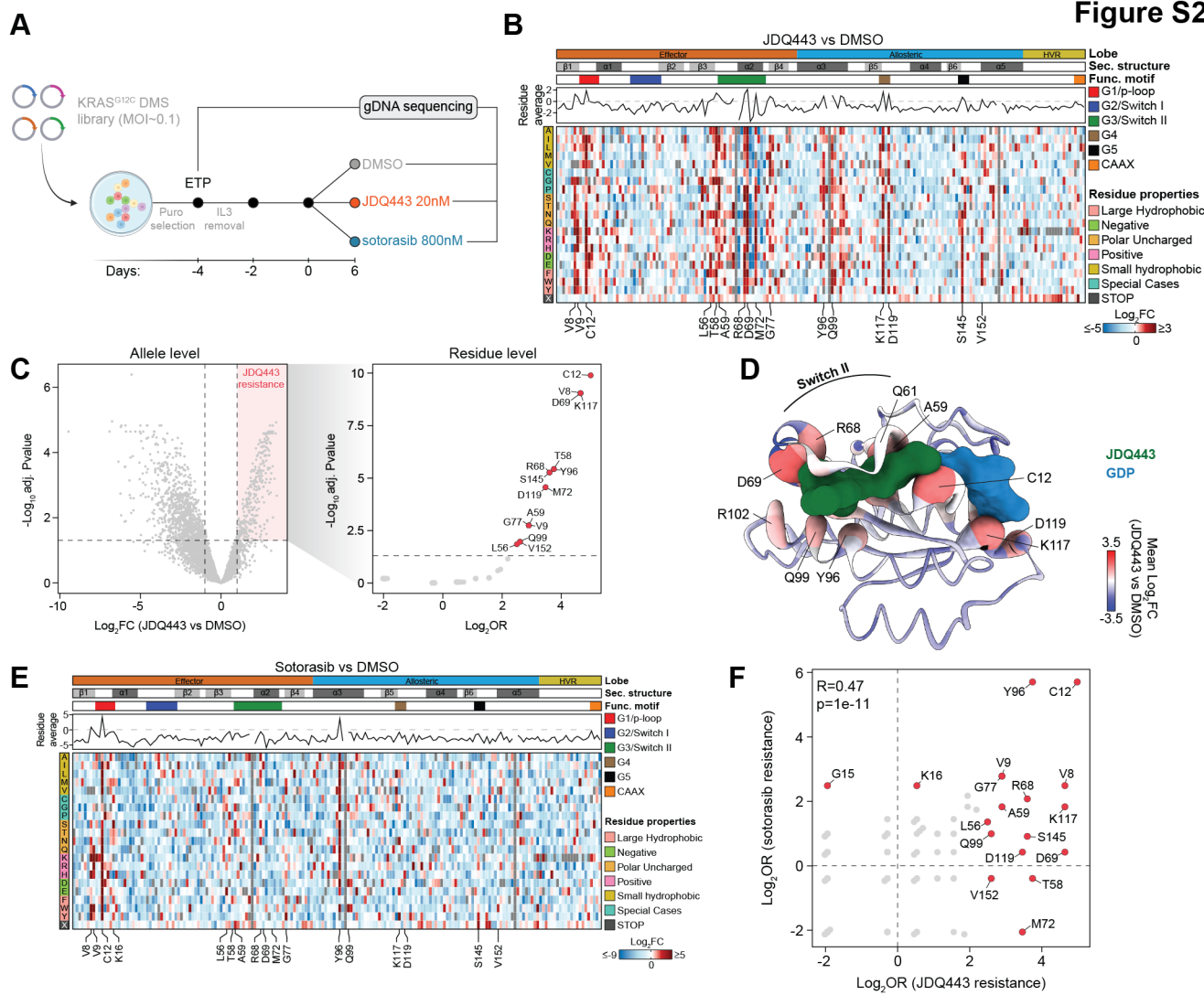

**Figure S3**

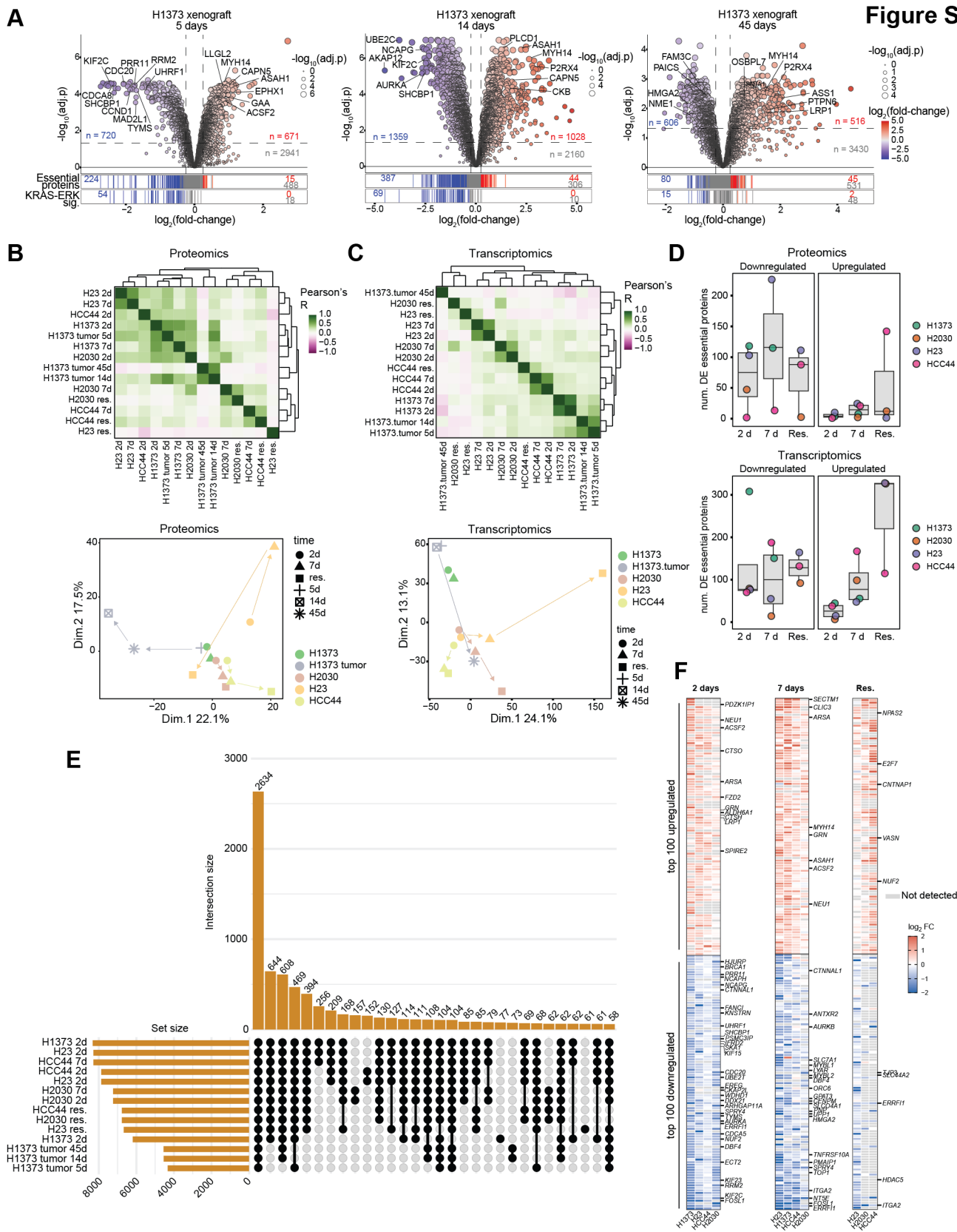

Figure S4

A

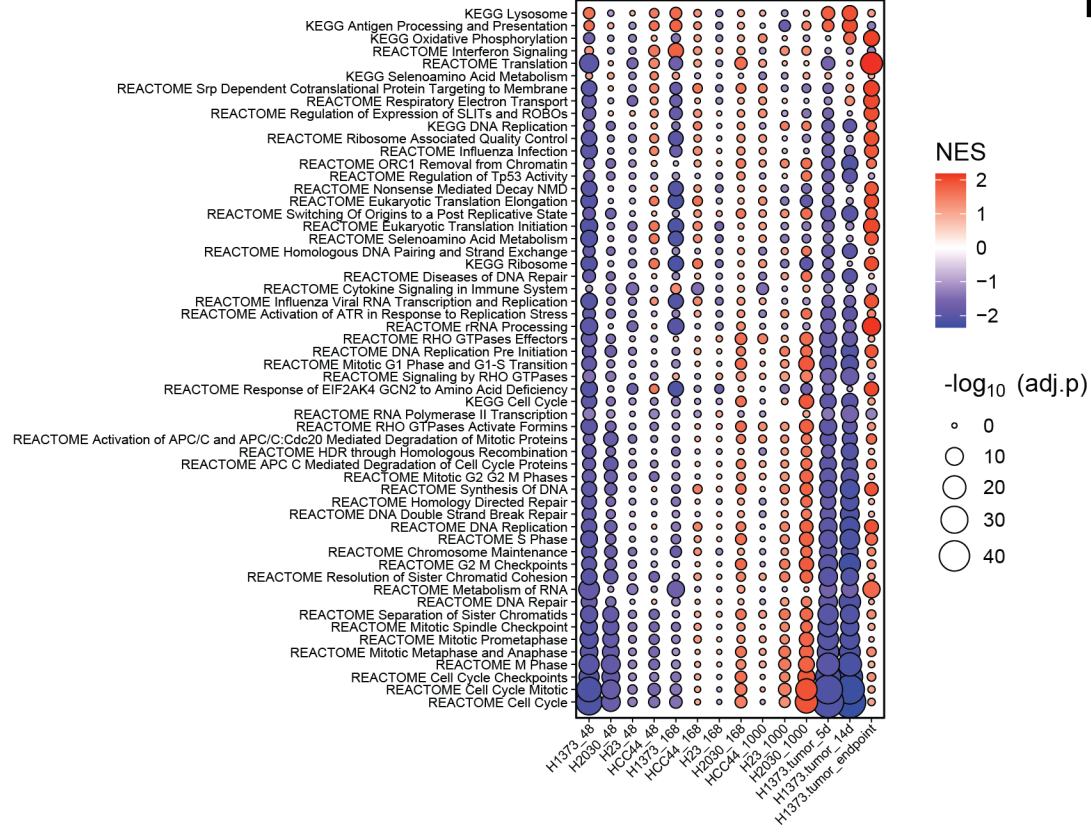

B

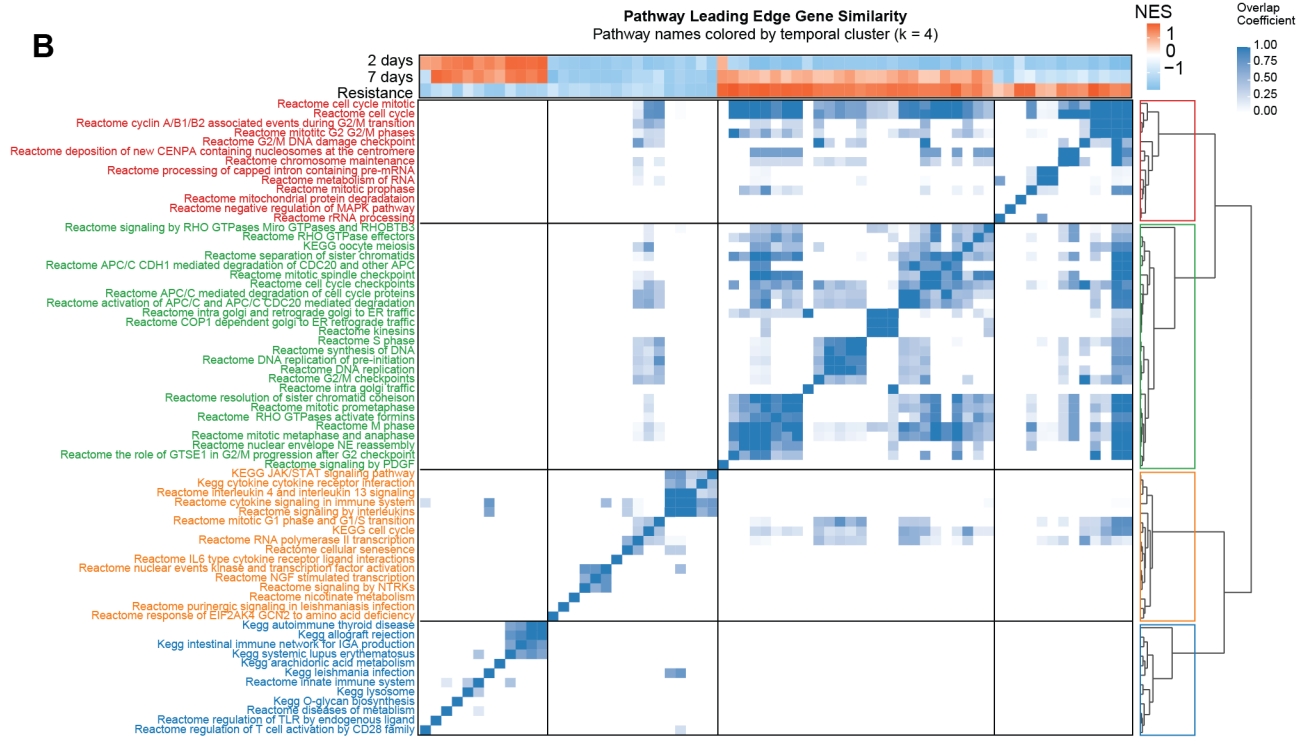

Figure S5

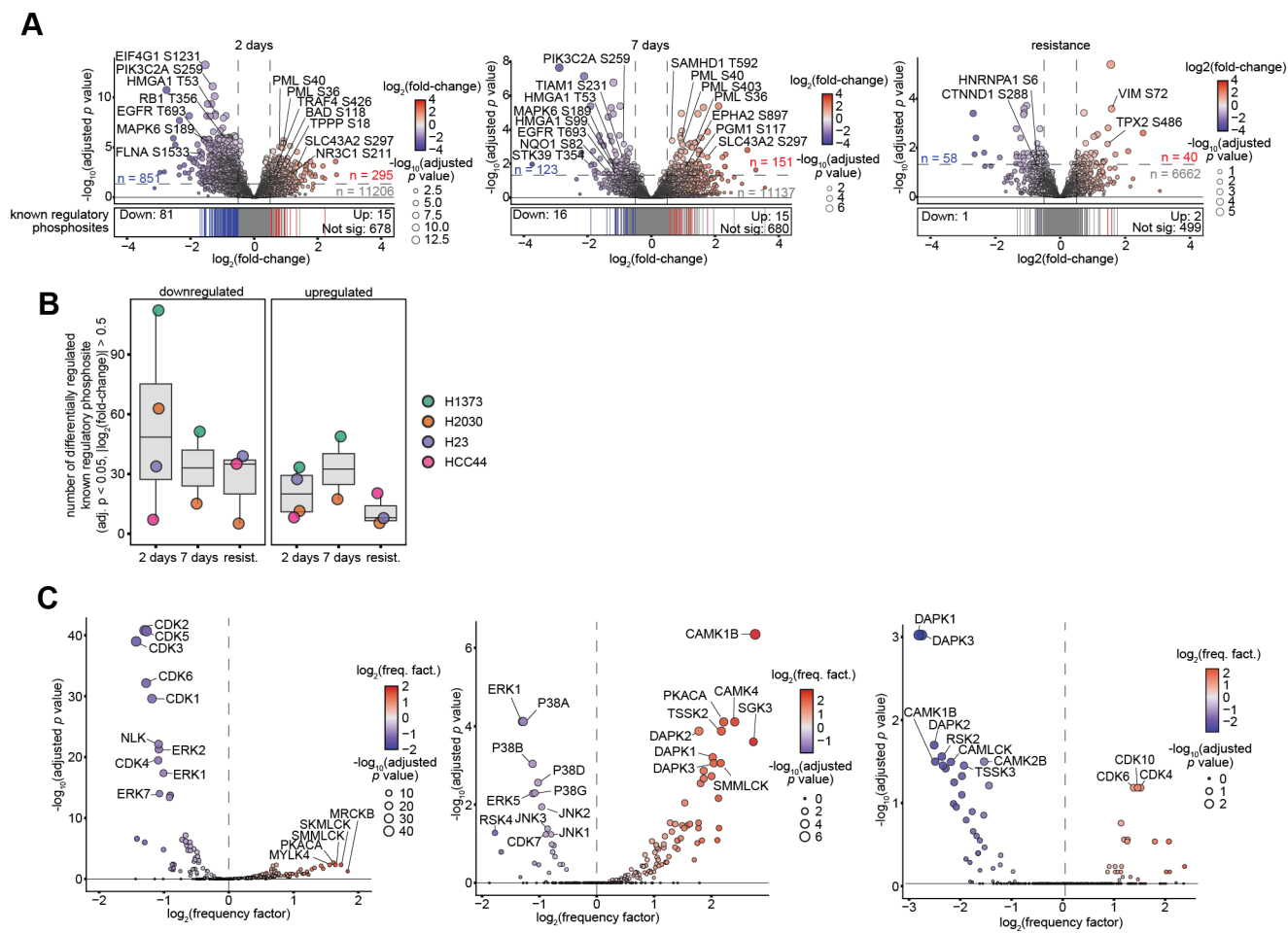

**Figure S6**

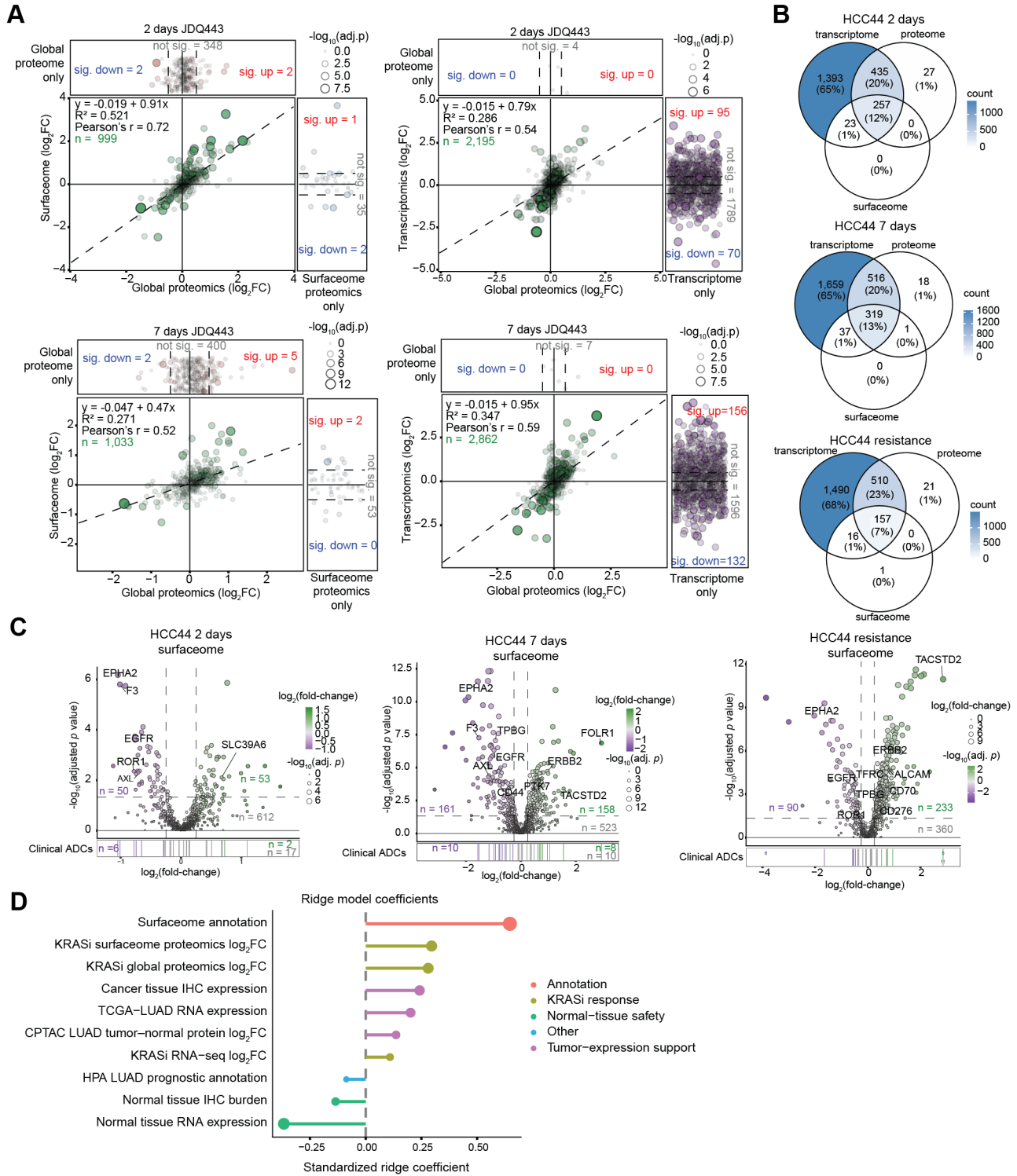

**FIGURE S7**

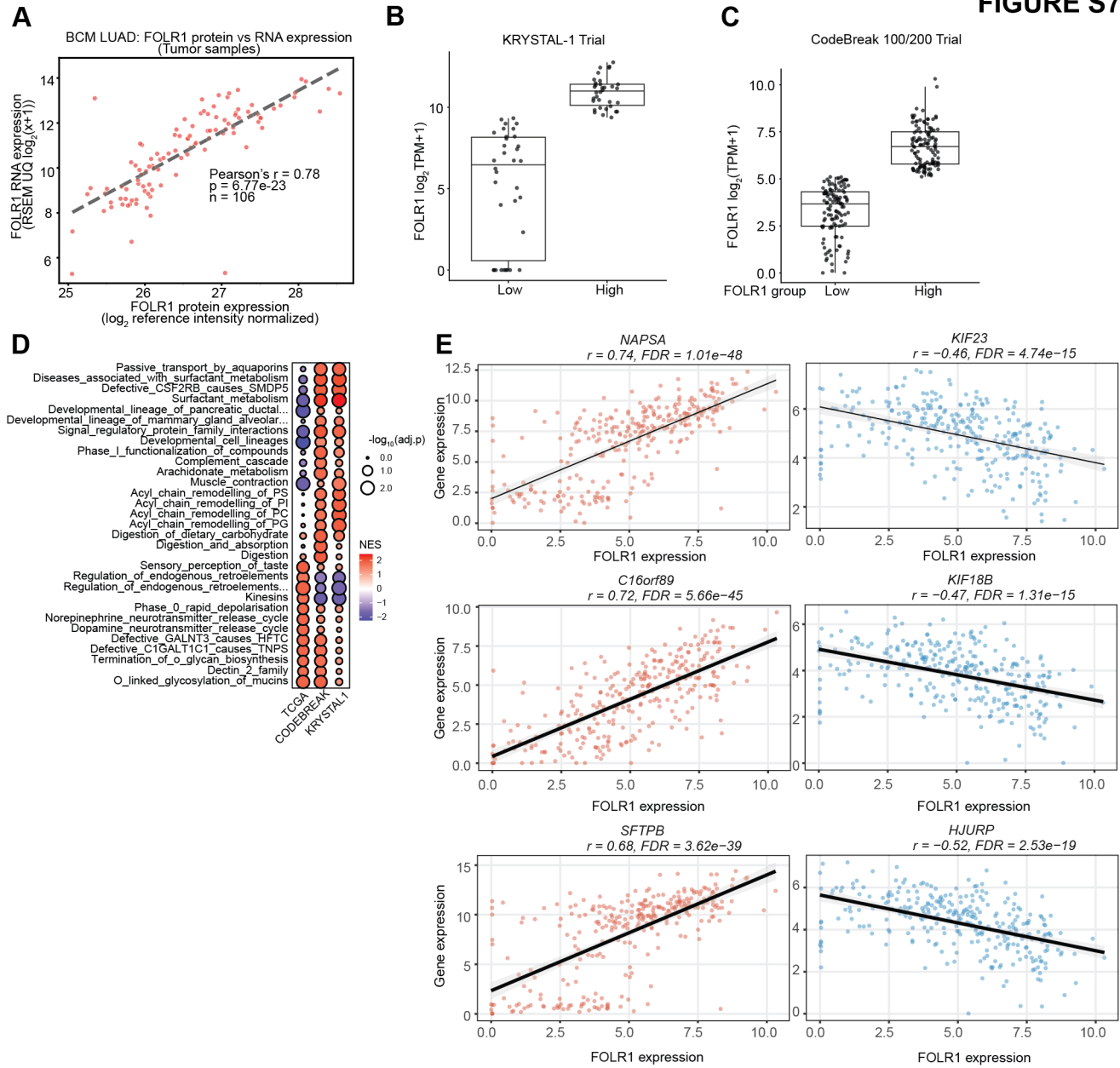

**Figure S8**

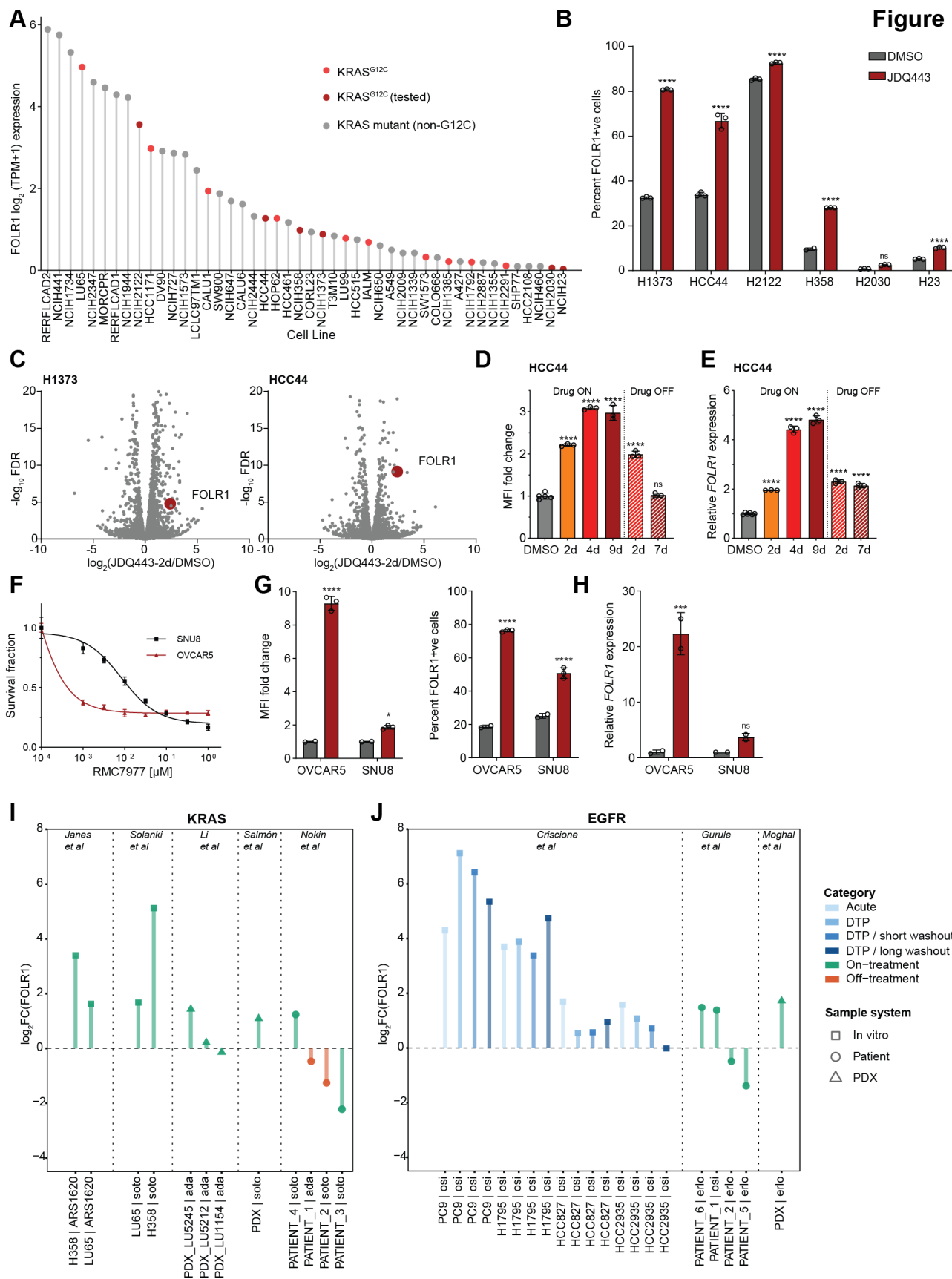

**Figure S9**

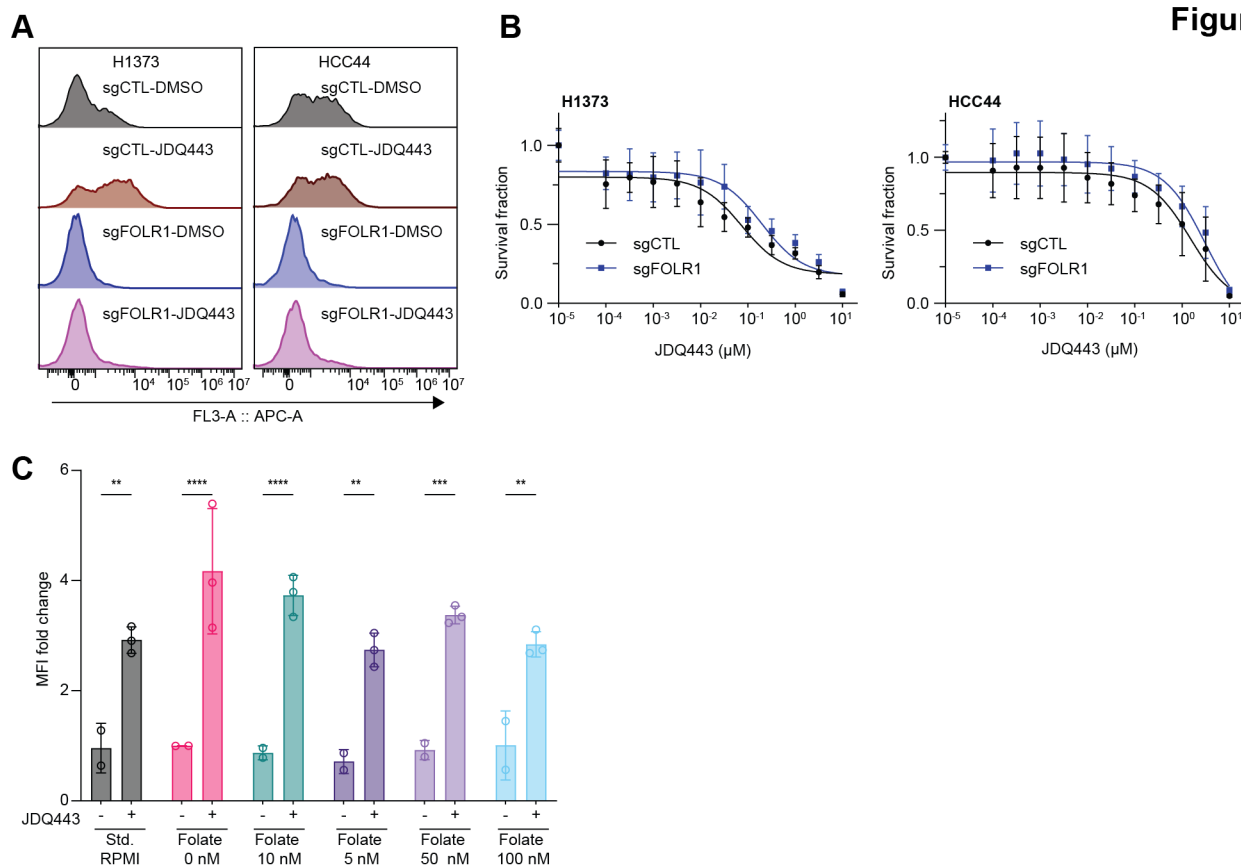

FIGURE S10

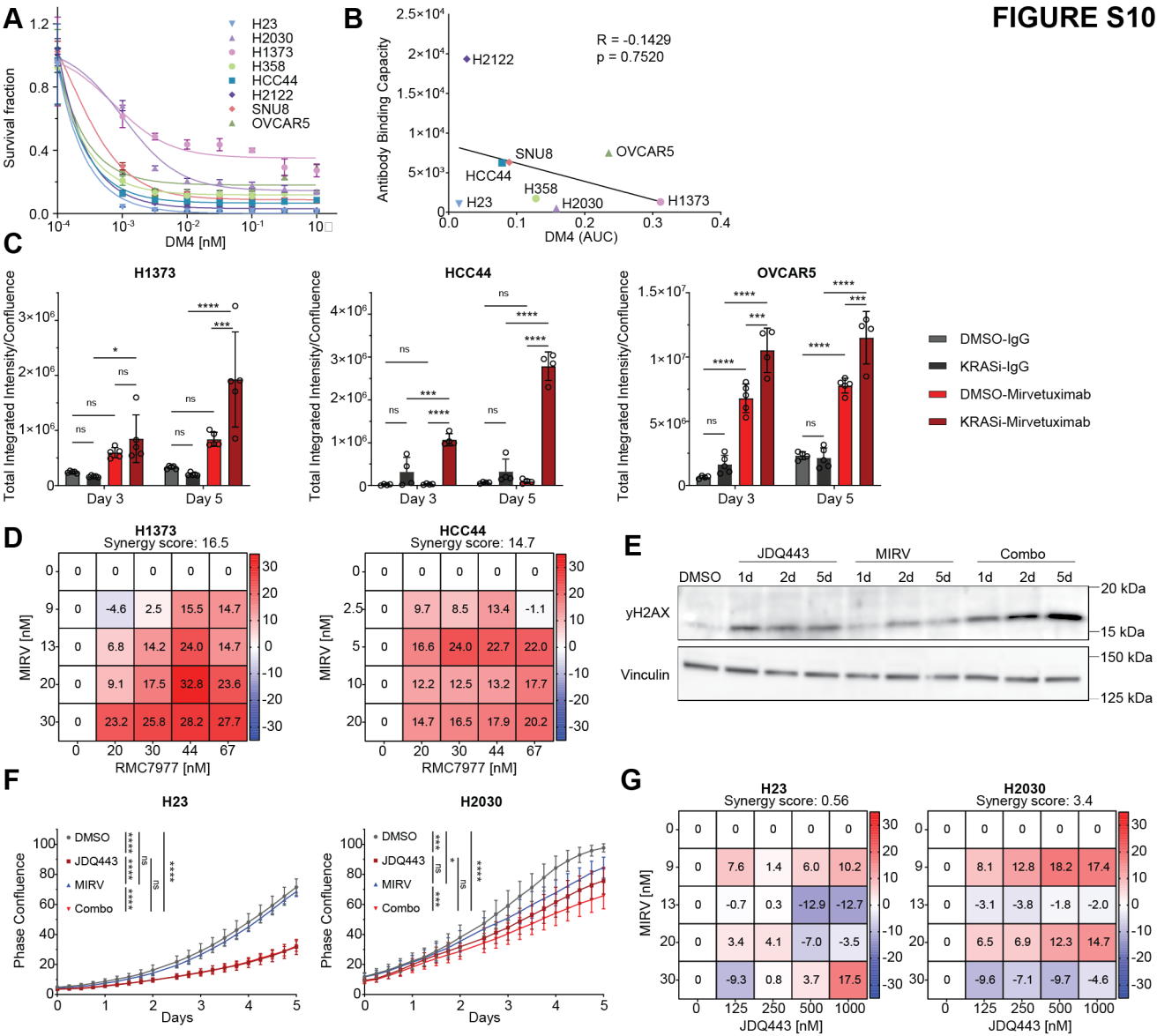

**FIGURE S11**

**A**

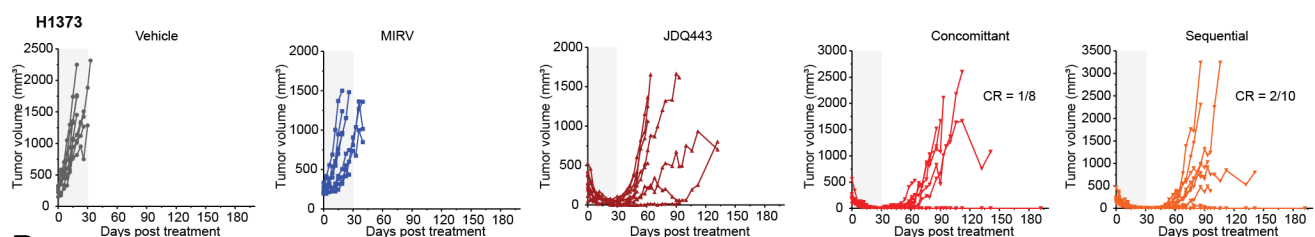

**B**

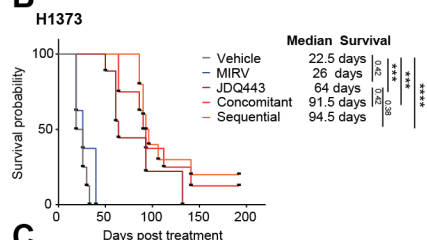

**C**

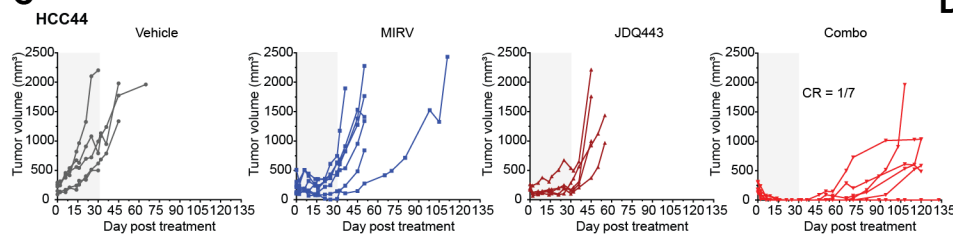

**D**

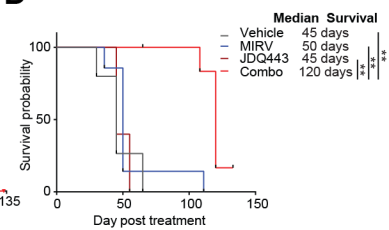

**E**

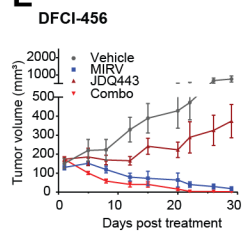

**F**

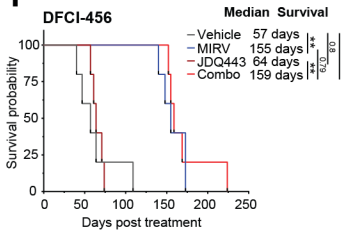

**G**

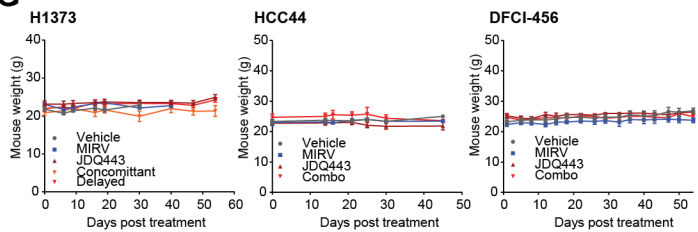
